# Structural and mechanistic basis for membrane recognition and activation of the vacuolar lipase Atg15

**DOI:** 10.64898/2025.12.23.696130

**Authors:** Tomoko Kawamata, Nobuo N Noda, Michiko Sasaki, Yoshinori Ohsumi, Yuji Sakai

## Abstract

Summary/Abstract

Atg15 is a vacuolar phospholipase B essential for the degradation of intravacuolar vesicles such as autophagic bodies. Despite its central role in cellular membrane turnover, the molecular basis of how Atg15 is activated and selectively acts on internal membranes has remained elusive. Here, by combining all-atom and coarse-grained molecular dynamics (MD) simulations with in vitro and in vivo analyses, we elucidate the structural and mechanistic principles underlying Atg15 activation and substrate recognition. Our simulations revealed that disulfide bonds are critical for maintaining the structural integrity of the catalytic core, while the C-terminal region locks the catalytic center in a closed state that prevents activation. Membrane binding induces a transition to an open state, enabling catalysis. Through MD-guided mutational analysis, we identified three regions crucial for catalytic locking, membrane binding, and substrate recognition, and experimentally confirmed that mutations in these regions inhibit activity. Furthermore, Atg15 preferentially associates with positively curved membranes, providing a potential basis for its selective action on internal vesicular membranes. These findings suggest that Atg15’s activity is controlled through multiple regulatory layers to ensure safe and selective membrane degradation.

## Introduction

Autophagy is an evolutionarily conserved degradation pathway in eukaryotes and enables cells to recycle cytoplasmic components through degradation in the lysosome or in the vacuole, in the case of yeast and plants. In yeast, a double-membrane autophagosome forms to sequester cytoplasmic material and deliver it to the vacuole. Upon fusion with the vacuolar membrane, the inner membrane structure—known as the autophagic body (AB)—is released into the vacuolar lumen, where it must be broken down to allow cargo degradation. Thus, degradation of the AB membrane represents an initial step of intravacuolar autophagic degradation. Failure to degrade the autophagic body leads to the accumulation of undegraded material and a loss of cell viability under nutrient starvation, resembling the phenotype observed in core autophagy gene mutants^1^.

The yeast vacuole contains a variety of hydrolases, such as proteases, nucleases, and lipases, responsible for degrading molecules delivered through the autophagic pathway^2–6^. Among them, Atg15 is a vacuolar protein harboring a lipase domain, and was initially identified in genetic screens as required for the degradation of autophagic bodies; deletion of *ATG15* causes the accumulation of autophagic bodies within the vacuole^5,6^. Atg15 can act on a broad range of intravacuolar membrane structures, including autophagic bodies, Cvt bodies, intraluminal vesicles, and potentially fragments of organellar membranes, such as those derived from the ER and mitochondria^6–9^. Atg15 consists of an N-terminal transmembrane domain (residues 13–35) and a lipase domain (residues 176–421) with the conserved GXSXG motif (residues 330–334), where S332 serves as the active-site residue (Fig. 1(**a**)). Atg15 is transported to the vacuole via the multivesicular body (MVB) pathway, a process that depends on its N-terminal region^10^.

**Figure 1.**
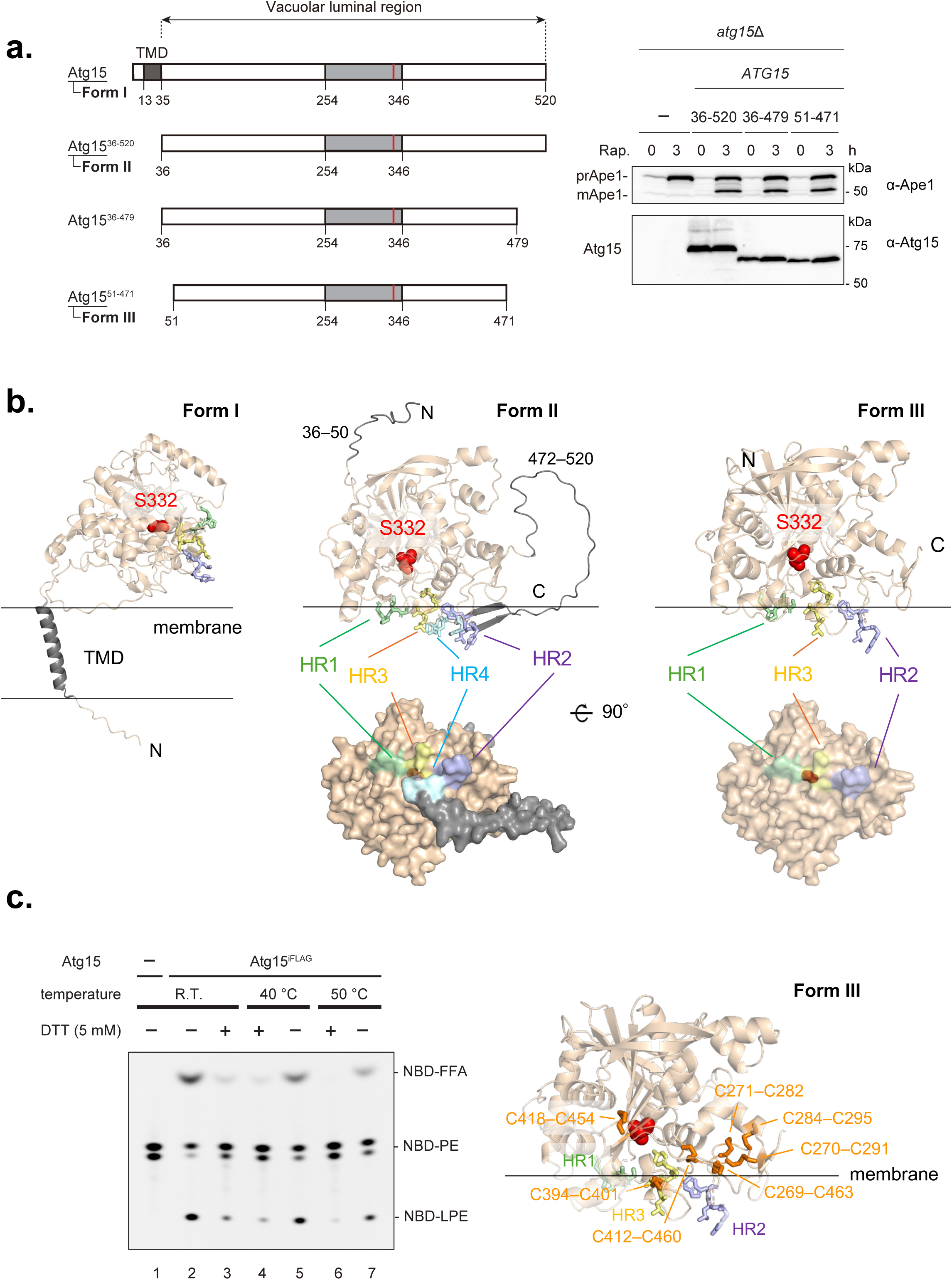
Structural and Functional Characterization of Atg15 Variants. (**a**) Schematic representation of Atg15 and CPY^N50^(vacuolar targeting signal)-fused Atg15 truncation derivatives, CPY^N50^–Atg15^36–520^, CPY^N50^–Atg15^36–479^, and CPY^N50^–Atg15^51–471^. The N-terminal transmembrane domain (TMD) and the conserved lipase consensus motif with Ser332 highlighted in red are shown. The functional activity of these constructs was analyzed by monitoring prApe1 maturation in *atg15*Δ cells. Cells expressing the indicated constructs under the endogenous *ATG15* promoter from a multicopy plasmid (pRS426) were grown in SD/CA medium and treated with rapamycin for 3 h. prApe1 and mApe1 were detected using α-Ape1 antibodies, and expression of Atg15 variants was confirmed by immunoblotting with α-Atg15 antibodies. **(b)** Predicted structures of the three Atg15 forms (Form I (the precursor, full-length protein), Form II (the N-terminally cleaved form, residues 36-520), and Form III (the C-terminally processed minimal functional region, residues 51–471)) —positioned relative to the membrane using the PPM server. The transmembrane domain (gray), the catalytic residue Ser332 (red) and membrane-interacting segments (HR1(246GLPG), pale-green; HR2(276YLW), light-blue; HR3(403LVGY), pale-yellow; HR4(511WLGF), pale-cyan) are highlighted. **(c)** (left) Effect of DTT on the lipase activity of Atg15. Purified Atg15^iFLAG11^ was pre-incubated with or without 5 mM DTT for 5 min at room temperature, 40°C, or 50°C. After cooling on ice, NBD-PE was added, and the lipase reactions were carried out at 30°C for 2 h. Lipids were then extracted, separated by TLC, and visualized by the fluorescence of NBD-PE. (right) AlphaFold2-predicted structure of Atg15 showing multiple disulfide bonds (orange) across different regions of the protein.

To clarify the biochemical basis of Atg15, we recently established a system to detect lipase activity within the yeast vacuole and demonstrated that Atg15 is the sole vacuolar lipase^11^. Previous studies revealed that its enzymatic activity strictly depends on the vacuolar proteases Pep4 and Prb1. Using vacuoles containing Pep4 and Prb1, we successfully isolated the active form of Atg15 and evaluated its activity. Atg15 has been shown to possess broad phospholipase B activity toward various phospholipid substrates^9,11^. Furthermore, the processed form of Atg15 alone was sufficient to disrupt purified ABs^8,11^.

However, several fundamental questions remain. First, it remains unclear why the vacuolar proteases Pep4 and Prb1 are essential for Atg15 function. These proteases are thought to mediate both the N-terminal cleavage of Atg15 upon its release from MVB vesicles and the subsequent processing at the C-terminal region within the vacuole^11^. However, the precise cleavage sites and their impact on the lipase activity of Atg15 remain unresolved. Second, although Atg15 hydrolyzes a wide spectrum of phospholipids, it does not act on the vacuolar limiting membrane. Our previous localization analysis showed that Atg15 preferentially associates with intravacuolar vesicles and is largely excluded from the vacuolar membrane, suggesting a mechanism of spatially restricted activation^11^. Third, the membrane recognition mechanism of Atg15 remains elusive. We have shown that Atg15 tightly associates with membranes even in the absence of its N-terminal transmembrane region^11^. However, it remains unknown how Atg15 detects specific features of membrane environments and how this recognition governs its membrane association. These unresolved issues are largely due to the challenges in purifying Atg15 in sufficient amounts and to its instability upon membrane dissociation. Atg15 is also a highly hydrophobic protein, which further complicates its biochemical handling. Although the structure of Atg15 has not been experimentally determined, an AlphaFold2-predicted model suggests that the catalytic center adopts a closed state. Molecular dynamics (MD) simulations also showed that Atg15 mainly adopts a closed state in solution, although an open state was observed in ∼ 5% of molecules upon cleavage between T153 and Y154^9^. How the transition to the open state is regulated, and particularly how the membrane association affects this closed-to-open transition, remains largely unknown.

Here, we combine all-atom MD simulations based on AlphaFold predictions with *in silico, in vitro, and in vivo* studies to elucidate the activation mechanisms of the lipase Atg15. We identify proteolytic processing and membrane association as key determinants of Atg15 activation and highlight four surface-exposed hydrophobic regions required for catalytic regulation, membrane binding and substrate engagement. We further show that positive membrane curvature promotes Atg15 membrane association and enhances lipase activity, suggesting a basis for its selective action on internal vesicular membranes.

## Results

### Defining the minimal essential region of Atg15 and structural insights into membrane interaction from AlphaFold2 predictions

We sought to define the minimal functional region of Atg15 from structural information and *in vivo* experiments as a basis for subsequent MD simulations. Our previous work showed that the N-terminal transmembrane region of Atg15 is dispensable and can be functionally replaced by a vacuolar targeting signal (Fig. 1(**a**))^11^. We showed that up to residue 58 could be deleted from the N terminus or up to residue 468 from the C terminus without loss of function. Because these analyses tested each terminus separately, we did not strictly define the minimal functional unit of Atg15. Closer inspection of the AlphaFold structural model revealed that several residues adjacent to these boundaries are likely important for structural stabilization. At the N terminus, F58 is buried in the hydrophobic core, and D51, K53, and T57 form hydrogen bonds or salt bridges with the main body of the protein (Supplementary Fig. S1). At the C terminus, residues 467–469 form part of a β-sheet, with F468 and I469 engaging in hydrophobic interactions and S471 forming a hydrogen bond, all of which appear to contribute to structural stability. Thus, we defined residues 51–471 as the structured region and tested its *in vivo* functionality using the Ape1 processing assay^11^. In this assay, precursor Ape1 (prApe1), a cargo of both the autophagy and Cvt pathways, is converted into the mature form (mApe1) upon Atg15-dependent disruption of autophagic bodies and Cvt bodies, followed by vacuolar protease processing. Since Ape1 maturation strictly depends on Atg15 activity, the prApe1/mApe1 ratio can be used to evaluate Atg15 function *in vivo*. Expression of this construct in *atg15*Δ cells complemented the defect in membrane degradation, and we therefore defined residues 51–471 as the minimal functional region of Atg15 (Fig. 1(**a**)). Given that proteolytic processing by the vacuolar proteases Pep4 and Prb1 is required for Atg15 function, we next addressed how Atg15 interacts with membranes at different forms of processing. To this end, we predicted the structures using AlphaFold2 and their orientations relative to the membrane using the PPM server^12^ for three forms of Atg15: Form I (the precursor, full-length protein), Form II (the N-terminally cleaved form, residues 36-520), and Form III (the C-terminally processed minimal functional region, residues 51–471).

Form I was predicted to have a hydrophobic helix spanning residues 11-35 inserted into the membrane (Fig. 1(**b**), left). The proteins in Forms II and III show a ∼90° rotation relative to Form I, bringing the membrane contact surface into contact with the membrane. Form II and III were predicted to have three hydrophobic regions—HR1 (246GLPG), HR2 (276YLW), and HR3 (403LVGY)— exposed on the surface, which interact with and insert into the membrane (Fig. 1(b) middle, right). In addition, Form II, but not Form III, was predicted to contain HR4 (511WLGF). In all these AlphaFold2 predicted structures, the active center, S332, was buried inside the protein and closed, inaccessible from the outside (Fig. 1(**b**)). The reason AlphaFold predicted only the closed state of Atg15 may be that it predicts the conformation assuming an aqueous environment and does not explicitly consider the membrane. In a later section, we will use MD simulations to analyze the dynamics and conformational change on the membrane due to interaction with lipids.

Structural prediction using AlphaFold2 indicated eight disulfide bonds in Atg15. To test their functional relevance, we examined the effect of conditions that break S-S bonds on Atg15 lipase activity. Purified Atg15 was incubated with or without DTT prior to the reaction with the substrate NBD-PE. DTT-treated Atg15 showed a marked reduction in lipase activity compared with untreated enzyme, although the activity was not completely abolished (Fig. 1(**c**) left). Because the reducing effect of DTT alone was insufficient at 30°C, we next combined DTT treatment with elevated temperatures (40°C and 50°C). While higher temperatures alone caused only a slight decrease in activity, the addition of DTT under these conditions almost completely abolished Atg15 activity. Furthermore, PPM analysis predicted that these disulfide bonds in Form III are localized near the active site at the membrane interface (between residues 269–295 and 401–460) (Fig. 1(**c**) right). Together, these results define residues 51-471 as the minimal functional region of Atg15, identify surface-exposed hydrophobic regions that mediate membrane association, and indicate that disulfide bonds proximal to the active site are important for Atg15 function.

### MD simulations of Atg15 highlight the role of the C-terminal region in regulating the open–closed transition

We next performed all-atom MD simulations to investigate how the C-terminal region and disulfide bonds influence the structural dynamics of Atg15. As a first step, we analyzed the conformational changes of Atg15 in aqueous solution without considering membrane lipid interactions. Three forms of Atg15 were simulated: Form III with disulfide bonds (Fig. 2(**a**, **b**) and Supplementary Fig. S2, Movie S1), Form II with disulfide bonds (Fig. 2(**c**, **d**) and Supplementary Figs. S3, S4, Movie S2), and Form III without disulfide bonds (Fig. 2(**e**–**g**) and Supplementary Fig. S5, Movie S3). In all cases, the initial structures were taken from the closed state predicted by AlphaFold2, in which the catalytic residue S332 is buried within the protein and inaccessible from the outside (Fig. 1(**b**)).

**Figure 2.**
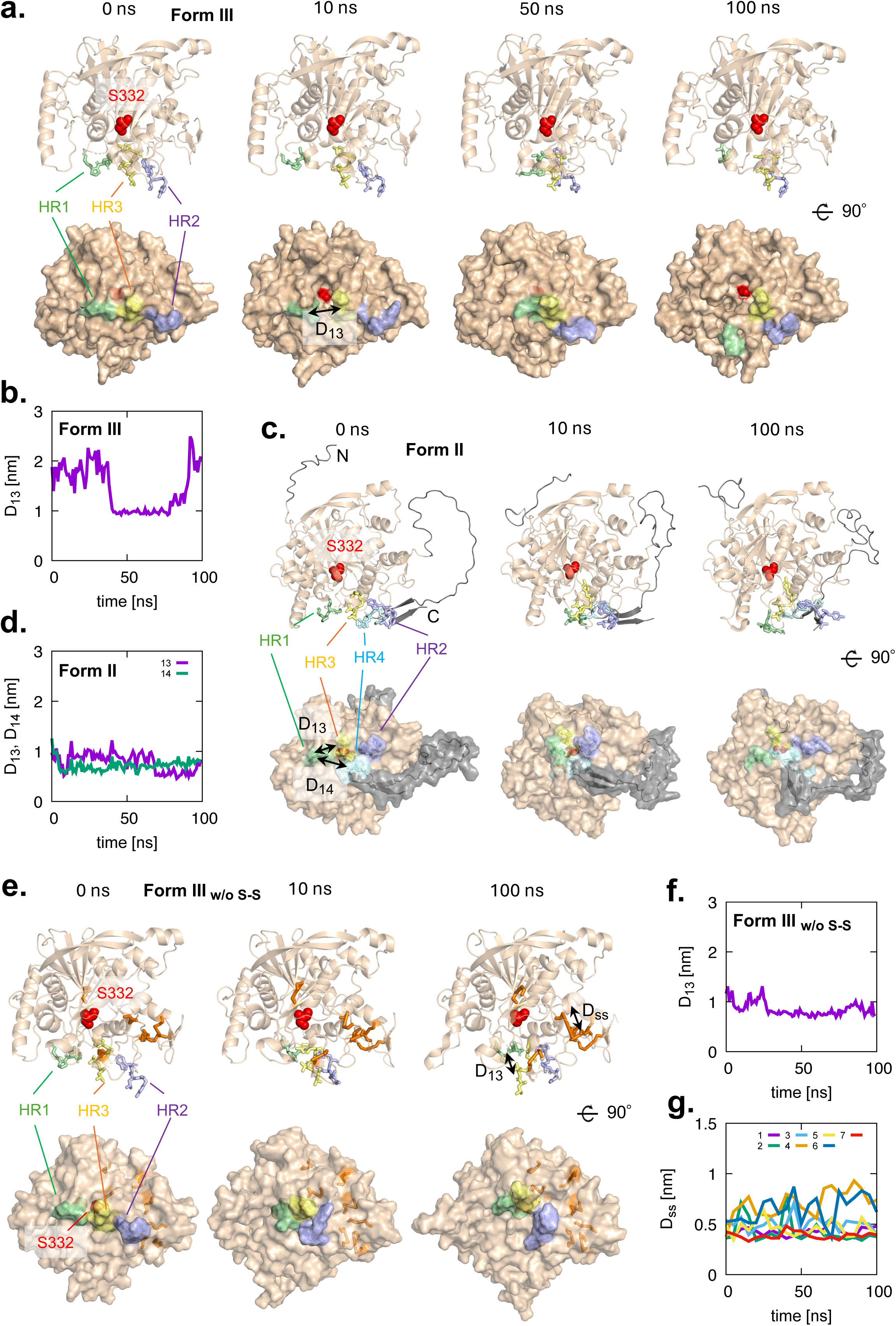
All-atom MD simulations of Atg15 in solution. (**a**, **b**) Form III (minimal functional region; residues 51–471) with disulfide bonds. (**c**, **d**) Form II (N-terminally cleaved form; residues 36–520) with disulfide bonds. (**e**–**g**) Form III without disulfide bonds. (**a**, **c**, **e**) Representative structures at 0, 10, 50, and 100 ns, respectively. Atg15, HR1, HR2, HR3, and HR4 are represented as wheat-brown, pale-green, light-blue, pale-yellow, and pale-cyan cartoons (top) and surfaces (bottom), respectively, while the catalytic residue Ser332 is represented as red spheres. (**b**, **f**) Time dependence of the distance between the centers of mass of HR1 and HR3, D_13_. (**d**) Time dependence of D_13_ and the distance between the centers of mass of HR1 and HR4, D_14_. (**g**) Time dependence of the distance between each sulfur atom forming disulfide bonds, D_ss_.

Simulations of Form III with disulfide bonds showed the interaction between the two hydrophobic regions exposed on the protein surface, HR1 and HR3, occasionally disengaging, resulting in the separation of the two hydrophobic sites and exposure of the active center S332 on the protein surface (Fig. 2(**a**) and Supplementary Fig. S2, Movie S1). This transition to the open state was transient, returning to the closed state in 50-60 ns. Six independent 100 ns simulations were performed from the same initial state; two remained closed, two immediately transitioned to the open state and remained open, and the remaining two transitioned between open and closed states at intervals of several tens of ns (Fig. 2(**b**) and Supplementary Fig. S2). These results indicate that Form III with disulfide bonds exists in equilibrium between the open and closed states, with comparable populations of both forms.

In contrast, simulations of Form II with disulfide bonds showed that the three hydrophobic regions exposed on the protein surface, HR1, HR3, and HR4, maintained tight hydrophobic interactions, with the active center remaining buried (Fig. 2(**c**) and Supplementary Fig. S3, Movie S2). The protein remained in the closed state for 100 ns in all five independent simulations (Fig. 2(**d**) and Supplementary Fig. S3) In this closed state, the C-terminal hydrophobic region (HR4; residues 511–514, WLGF) was inserted between residues 392–411 (HR3) and 241–258 (HR1), thereby locking the structure in the closed state (Supplementary Fig. S4). L512 formed direct hydrophobic interactions with P248, V404, and F433, stabilizing the autoinhibitory state. In addition, F514 interacted with the underlying helix, further contributing to the maintenance of the closed state. Altogether, these results suggest that the C-terminal region of Atg15, especially HR4, functions as an autoinhibitory region and negatively regulates the transition to the open state by locking the HR1–HR3 interaction, thereby reinforcing the closed state of Atg15.

Furthermore, in simulations of Form III in which disulfide bonds were removed to model the reduced (–SH) state, the hydrophobic interaction between HR1 and HR3 was stably maintained for 100 ns, with no active center exposed on the protein surface in all five independent simulations (Fig. 2(**e, f**) and Supplementary Fig. S5. Movie S3). The S–S distances of the pairs that originally formed disulfide bonds increased (Fig. 2(**g**)) and the overall structure became disrupted in some cases. Notably, all disulfide bonds are positioned near HR3, on the opposite side of HR1 (Fig. 1(**c**), right), suggesting that they stabilize the structure around HR3 and thereby preventing excessive HR1–HR3 association. These results suggest that disulfide bonds inhibit excessive proximity between HR1 and HR3 via hydrophobic interactions, thereby facilitating the transition of Atg15 to the open state. This interpretation is consistent with our experimental observation that reduction of disulfide bonds by DTT treatment abolished lipase activity (Fig. 1(**c**), left). In summary, our MD simulations show that the C-terminal region (HR4) locks Atg15 in a closed state and suppresses the closed-to-open transition, whereas disulfide bonds promote the closed-to-open transition by permitting HR1–HR3 disengagement and exposure of the catalytic site.

### Membrane interactions stabilize the open state of Atg15 and promote lipid access to the catalytic center

MD simulations in aqueous solution showed that the transient interaction between two surface-exposed hydrophobic regions, HR1 and HR3, is critical for the conformational transition to the active state, but this transition is suppressed when HR4 is present. This raised the question of how the open–closed transition of Atg15 would behave when simulated in the presence of a membrane. Our previous work demonstrated that Atg15 tightly associates with membranes even without its N-terminal transmembrane region^11^. Consistently, structural prediction using the PPM server indicated that several hydrophobic regions (HR1–HR3) insert into the membrane (Fig. 1(**b**), right). Here, we used MD simulations to investigate how the interaction of these exposed hydrophobic regions with the membrane affects the transition to the active state and the approach and entry of a single membrane phospholipid substrate into the catalytic center S332.

First, we investigated the effect of membrane binding on Form III with disulfide bonds (Fig. 3(**a**) and Supplementary Figs. S6, S7, Movie S4). All membrane simulations were performed using a pure POPC bilayer as a model membrane. In the initial configuration, three hydrophobic regions interacted with the membrane, with HR2 and HR3 strongly and HR1 weakly binding to the membrane. After a few tens of nanoseconds, HR1 detached from the membrane, leading to a transition to the open state with the catalytic center S332 exposed. HR2 and HR3 remained inserted into the membrane, and once released, HR1 did not rebind, leaving Atg15 stably associated with the membrane in the open state. The transition to the open state occurred within 20 ns and persisted throughout the subsequent 100 ns in all five independent simulations (Fig. 3(**b**) and Supplementary Fig. S6).

**Figure 3.**
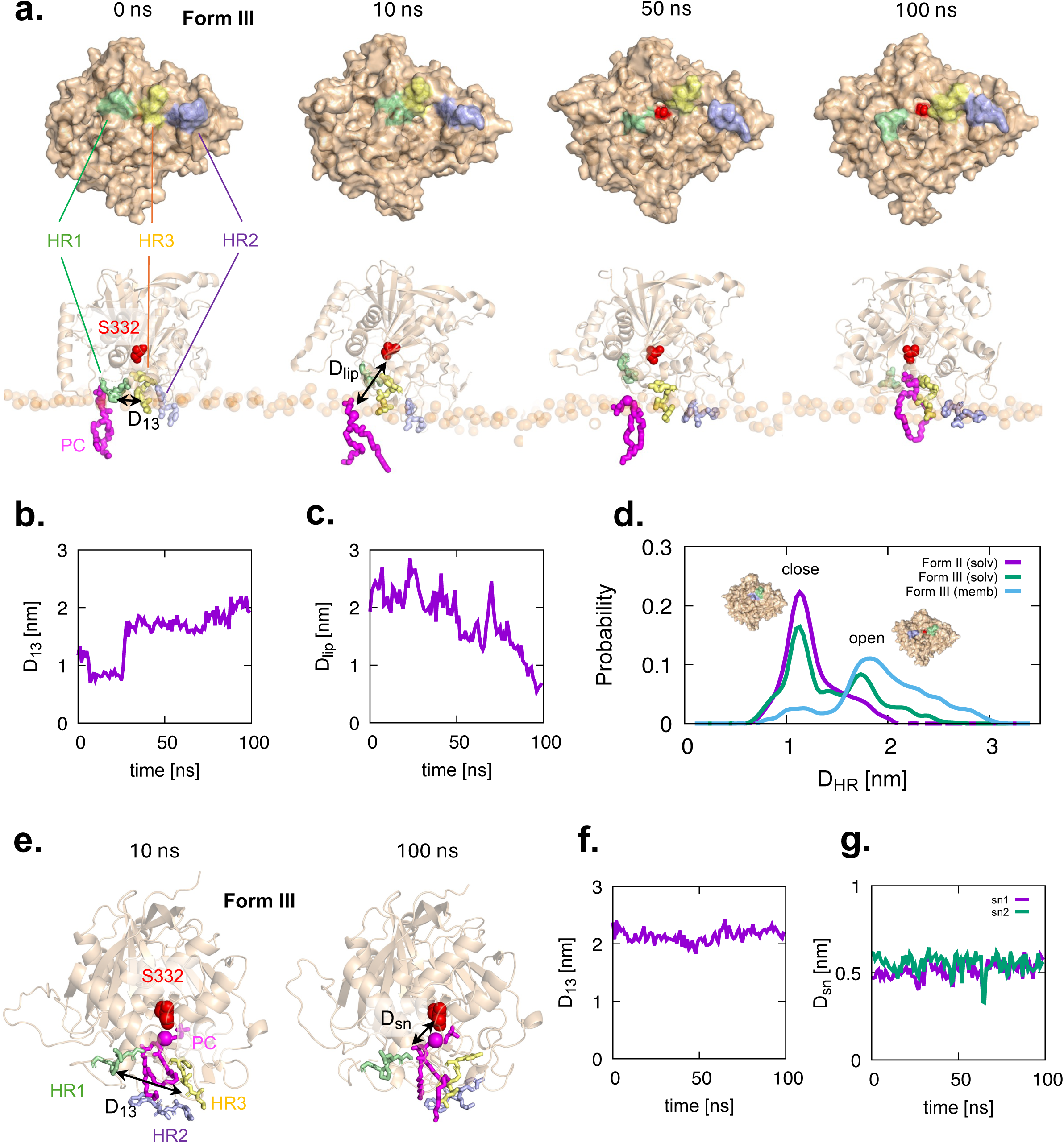
All-atom MD simulations of Form III with disulfide bonds in different POPC environments. (**a**–**d**) On a POPC bilayer. (**a**) Representative structures at 0, 10, 50, and 100 ns, respectively. Each representation is the same as in Fig. 2(a). The lower panels illustrate a POPC molecule (magenta sticks) being captured from a POPC bilayer (phosphorus as pale orange spheres) and guided toward the catalytic center. (**b**) Time dependence of the distance D_13_. (**c**) Time dependence of the distance between the nucleophile (oxygen of the hydroxyl group) of residue S332 and the phosphorus of a lifted phospholipid molecule, D_lip_. (**d**) Probability distributions of the open and closed states of Atg15: Form III in solution, on the POPC membrane, and Form II in solution. (**e**–**g**) With a single POPC molecule. (**e**) Representative structures at 10 and 100 ns, respectively. Each representation is the same as in panel (**a**) bottom. (**f**) Time dependence of the distance D_13_. (**g**) Time dependence of the distances between the nucleophile (oxygen of the hydroxyl group) of residue S332 and the carbonyl oxygens at the *sn-1* and *sn-2* positions of the phospholipid, D_sn_.

In addition, during the 100 ns simulation, a POPC molecule located directly beneath Atg15 was occasionally lifted from the lipid bilayer, approaching the catalytic center (bottom panel of Fig. 3(**a**) and Fig. 3(**c**)). The pulled lipid approached to within ∼1 nm of the catalytic center, and in three out of five independent 100 ns simulations, such lipid lifting events were observed (Supplementary Fig. S7). Fig. 3(**d**) shows the probability distributions of the open and closed states of Atg15 in solution and on the membrane. The minimal functional region exhibited a slightly open state in solution, whereas on the membrane the open state was predominantly favored. In summary, the minimal functional region opened more readily on the membrane, stabilizing its catalytic center in the open state.

Second, we further performed simulations of the minimal functional region in the presence of a single POPC molecule (Fig. 3(**e**) and Supplementary Fig. S9, Movie S5). During the 100 ns simulation, the distance between residues HR1 and HR3, as well as the distance between the hydroxyl group of S332 and the carbonyl carbons of both the *sn-1* and *sn-2* positions of POPC, remained largely unchanged (Fig. 3(**f**, **g**)). This orientation would allow potential attacks on both *sn-1* and *sn-2* ester bonds, which is consistent with the phospholipase B activity of Atg15. The combined hydrophobic surfaces formed by residues HR1, HR2, and HR3 appeared to act as a trap for POPC, likely contributing to the retention of the acyl chains. In summary, membrane binding stabilizes the open conformation of Atg15 and enables substrate capture and positioning through the combined contribution of HR1–HR3, retaining the substrate’s acyl chains near the catalytic center.

### Coordinated roles of HR1–HR3 in conformational regulation, membrane binding, and substrate recognition of Atg15

In the previous section, MD simulations showed that the hydrophobic regions HR1, HR2 and HR3 promote membrane binding, thereby facilitating the transition to the open state and enabling phospholipid entry into the catalytic center. To experimentally evaluate the importance of these three hydrophobic regions, we next examined how converting each hydrophobic region (HR1**–**HR3) into a negatively charged segment affects the lipase activity of Atg15. We introduced triple or quadruple Asp substitutions (HR1D mutant: 246–249 (GLPG → DDDD), HR2D mutant: 276–278 (YLW → DDD), and HR3D mutant: 403–406 (LVGY → DDDD)). This design allowed us to assess *in vivo* how disruption of locking or membrane association affects Atg15 function. We first examined the activity of these mutants using the Ape1 processing assay, which serves as an *in vivo* readout of Atg15 lipase activity toward native membrane substrates (ABs). Mutations in the hydrophobic regions abolished or reduced Atg15 activity: the HR2D mutant (Y276D/L277D/W278D) and the HR3D mutant (L403D/V404D/G405D/Y406D) completely lost activity, whereas the HR1D mutant (G246D/L247D/P248D/G249D) retained partial activity (Fig. 4(**a**)). In parallel with the experiments, we analyzed the interactions of these mutants with the membrane using MD simulations. As the initial state, each mutant was placed on the membrane with the same orientation and position as the wild-type predicted by PPM. Under these conditions, the HR1D mutant remained membrane-associated, whereas HR2D and HR3D dissociated from the membrane within 100 ns (Fig. 4(**b**, **d**, **f**) and Supplementary Figs. S10–S12). Furthermore, the HR1D mutant opened its active site (Fig. 4(**c**) and Supplementary Fig. S10), whereas the HR2D and HR3D mutants failed to open (Fig. 4(**e**, **g**) and Supplementary Figs. S11, S12). These findings demonstrate that the exposed hydrophobic regions, especially HR2 and HR3, are essential for Atg15 membrane association, and inhibition of this association impairs its lipase function.

**Figure 4.**
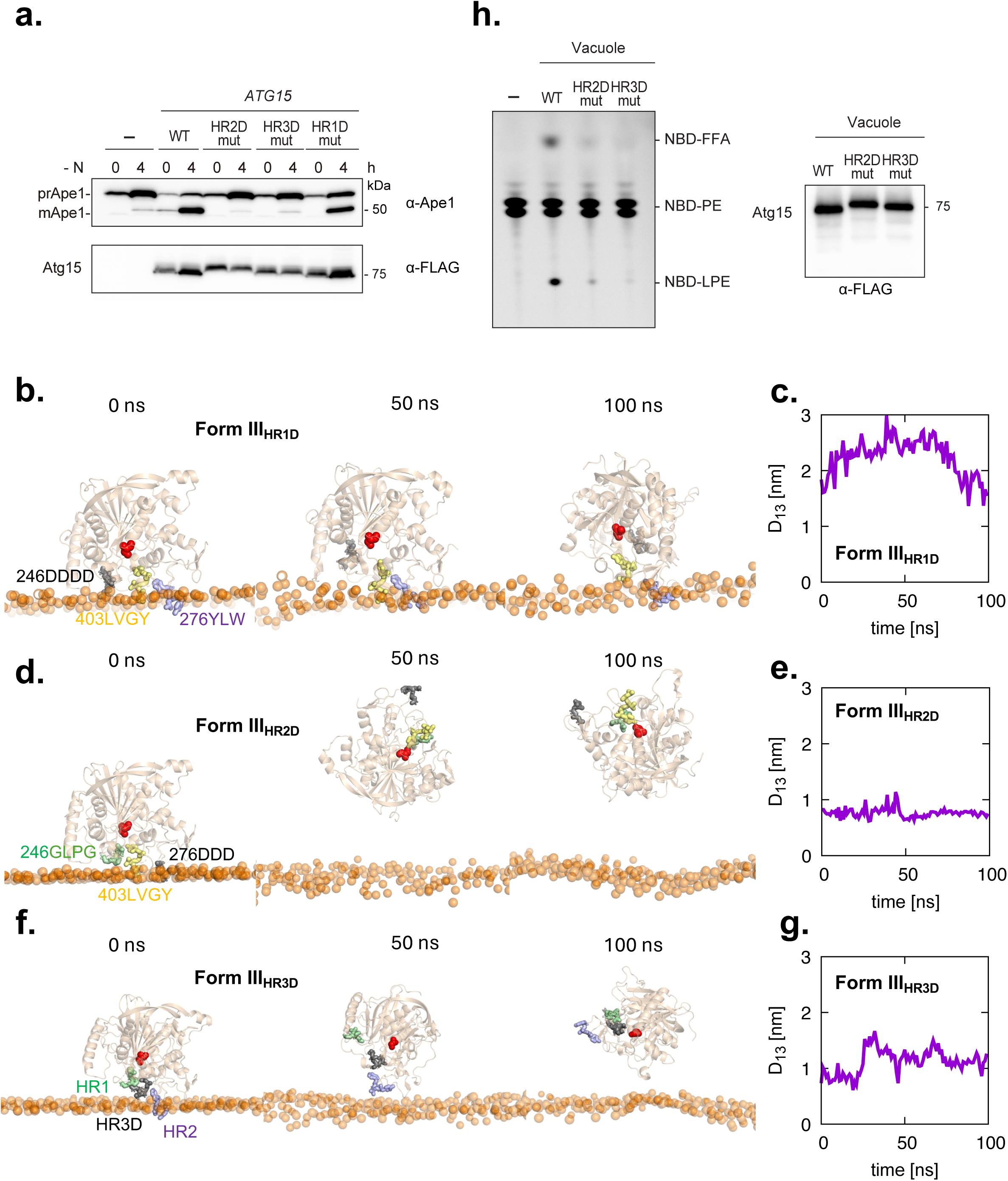
Experimental and simulation analysis of Atg15 mutants. (**a**) Maturation of prApe1 and expression of internally FLAG-tagged Atg15 and its HR1D, HR2D, and HR3D mutants in *atg15*Δ cells. Cells were grown in SD/CA medium or shifted to SD-N medium for 4 h. prApe1 and mApe1 were detected using anti-Ape1 antibodies, and Atg15 was detected using anti-FLAG antibodies. (**b**, **d**, **f**) All-atom MD simulations of Atg15 mutants on a POPC membrane. Representative structures at 0, 50, and 100 ns for the HR1D, HR2D, and HR3D mutants, respectively. Each representation is the same as in Fig. 2(a), but the mutated regions are shown in black sticks. (**c**, **e**, **g**) Time dependence of the distance D_13_ for the corresponding mutants. (**h**) Lipase activity of vacuolar lysates from *atg15*Δ cells harboring *ATG15*, *ATG15^HR2D^*or *Atg15^HR3D^* (left). Expression levels of Atg15 in each strain are shown (right).

Building on our *in vivo* assays showing that HR2D and HR3D are largely nonfunctional whereas HR1D displays a partial defect with native bilayer substrates (ABs) (Fig. 4(**a**)), we next examined whether these regions support lipase activity toward a non-membranous substrate. Using NBD-PE as a free phospholipid substrate, HR2D and HR3D showed no detectable hydrolysis compared with the wild type (Fig. 4(**h**)), indicating that both motifs are required for phospholipid recognition. Integrating these results with our MD simulations, we conclude that the three hydrophobic regions act in a coordinated manner: HR3 is essential for catalytic-center locking (Figs. 2, 3), membrane association (Fig. 3(**a**) and Fig. 4(**a**)), and substrate recognition (Fig. 3(**e**) and Fig. 4(**h**)); HR2 is likewise indispensable, contributing primarily to membrane binding (Fig. 3(**a**) and Fig. 4(**a**)) and substrate recognition (Fig. 3(**e**) and Fig. 4(**h**)); and HR1 is required for full activity and likely supports catalytic-center locking (Figs. 2, 3) and substrate recognition (Fig. 3(**e**)) once Atg15 is membrane-associated.

### Positive membrane curvature promotes Atg15**–**membrane interaction and enhances lipase activity

We next examined whether the lipase activity of Atg15 is influenced by membrane curvature. In our previous study, we established a liposome-based NBD-PE degradation assay and showed that Atg15 can degrade NBD-PE in POPC-based liposomes^11^. We further demonstrated that Atg15 was sufficient to disrupt autophagic bodies (ABs, ∼400 nm in diameter) purified in vitro. Building on these findings, we next prepared liposomes of various sizes composed of POPC (94%) and NBD-PC (6%) to systematically investigate the influence of membrane curvature on Atg15 activity. The hydrated lipid suspension was sequentially extruded through polycarbonate membranes with pore sizes of 1000, 400, 100, and 50 nm to generate liposomes of identical composition but different sizes (Supplementary Fig. S13(**a**)). Purified Atg15 was incubated with these liposomes, and the cleavage of NBD-PC was quantified. The results revealed that degradation of NBD-PC was more efficient in 50 nm and 100 nm liposomes than in 400 nm or 1000 nm liposomes (Fig. 5(**a**)). To confirm that this effect was not specific to the lipid species, we repeated the experiment using liposomes containing NBD-PE instead of NBD-PC. A similar trend was observed (Supplementary Fig. S13(**b**, **c**)). These findings indicate that Atg15 lipase activity is enhanced on membranes with higher curvature.

**Figure 5.**
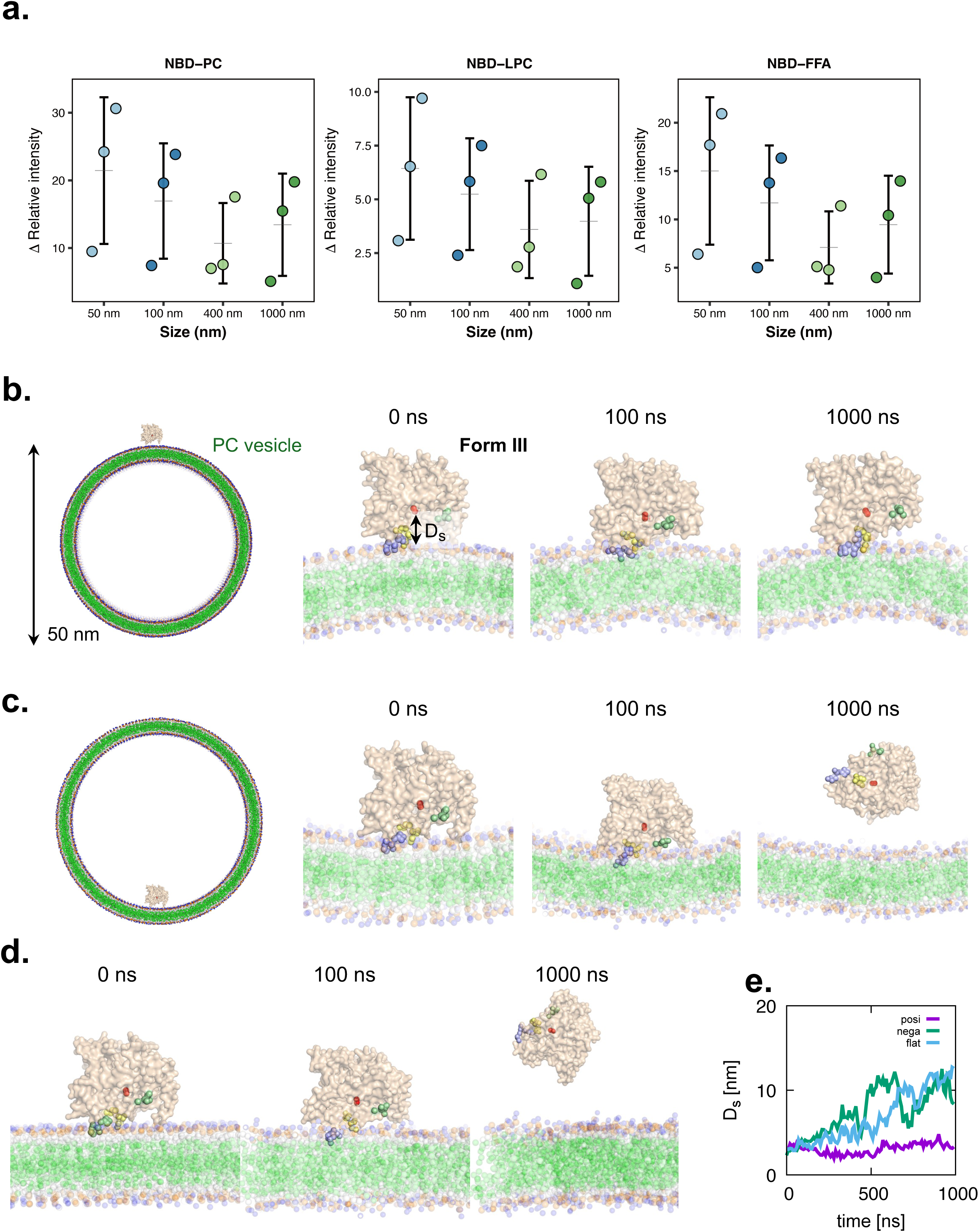
Membrane curvature modulates the lipase activity of Atg15. (**a**) Lipase activity of purified Atg15 toward liposomes prepared by extrusion through filters of different pore sizes. POPC liposomes containing 6% NBD-PC were prepared by sequential extrusion through polycarbonate filters with nominal pore sizes of 1000, 400, 100, and 50 nm. The liposomes were incubated with purified wild-type Atg15 or Atg15(S332A), and the reactions were analyzed by TLC. The amounts of NBD-PC, NBD-LPC, and NBD-FFA were quantified. The relative proportions of each lipid species were calculated, and the differences between wild-type Atg15 and Atg15(S332A) were expressed as ΔRelative intensity. The actual liposome diameters determined by DLS are provided in Supplementary Fig. S13(**a**). (**b**–**d**) Coarse-grained MD simulations of the Atg15 Form III on membranes with different curvatures. Simulation results of representative structures at 0, 100, and 1000 ns for Atg15 initially positioned on the outer surface with positive curvature (**b**), inner surface with negative curvature (**c**) of a POPC vesicle with 50 nm diameter and positioned on a flat POPC bilayer (**d**). Atg15, HR1, HR2, HR3, catalytic residue S332, the POPC tail are represented by wheat-brown surface, pale-green, light-blue, pale-yellow, red, and green spheres, respectively. (**e**) Time dependence of the distance between the center of mass of S332 and the nearest POPC phosphate group, D_s_.

To understand the curvature dependence of Atg15 lipase activity at the molecular level, we performed MD simulations of Atg15 on membranes with different curvatures. To be consistent with experimental conditions, the simulations were performed with Atg15 positioned on the outer surface of a 50 nm diameter POPC vesicle with positive curvature. As controls, similar simulations were also performed with Atg15 positioned on a flat POPC membrane surface and on the inner surface of a 50 nm diameter POPC vesicle with negative curvature, respectively. Because of the large system size, we performed coarse-grained simulations using the MARTINI model. The MD simulations revealed that Atg15 stably maintained its association with positively curved membranes for 1 µs, whereas it tends to dissociate from negatively curved or flat membranes within several hundred nanoseconds (Fig. 5(**b**–**e**), Supplementary Fig. S14). Together, the results from the liposome assay and simulations suggest that Atg15 preferentially binds to highly positively curved membranes, thereby enhancing its lipase activity.

## Discussion

Our simulations uncovered several key aspects of the conformational dynamics of Atg15, offering insights that extend beyond what could be achieved by biochemical assays alone. Both biochemical analyses and simulations demonstrated that the disulfide bonds are critical for maintaining the proper structure and function of Atg15 (Figs. 1(**c**), 2(**e**–**g**)). Given that the vacuolar lumen represents an oxidative environment, it is reasonable that Atg15 can function while undergoing closed-to-open conformational transitions with disulfide bonds intact. In our simulations, the catalytic site of Atg15 occasionally adopted an open state in solution, but this state was unstable (Fig. 2(**a**, **b**)). By contrast, upon association with membranes, the catalytic center adopted and maintained a stable open state, thereby permitting substrate access and catalysis (Fig. 3(**a**–**d**)). These findings suggest that membrane association triggers and stabilizes the open state of the catalytic center, thereby enabling Atg15 activity. In other words, Atg15 appears to be intrinsically coupled to the membrane environment, with its catalytic competence being gated by membrane association.

Building on this, our analyses also provided new insights into how specific hydrophobic regions regulate Atg15 activity (Figs. 3, 4). HR1 and HR3 work in concert, enforcing autoinhibition of the catalytic center while facilitating retention of substrate lipid acyl chains, as underscored by our MD simulations. These two regions appear to mediate only partial autoinhibition in solution, limiting premature activation of the enzyme, whereas the additional contribution of the C-terminal region (HR4) establishes a more rigid lock that fully reinforces the autoinhibited state (Fig. 2(**c**, **d**)). In the absence of HR4, the inhibitory interaction between HR1 and HR3 is released upon membrane binding, thereby allowing activation of the catalytic site (Fig. 3(**a**–**c**)). This finding provides a mechanistic explanation for why proteolytic cleavage of the C terminus by Pep4/Prb1 is essential for Atg15 function, as removal of the C terminus relieves the inhibitory lock on the catalytic center. This model is also consistent with our truncation experiments showing that the C terminus is dispensable for enzymatic activity (Fig. 1(**a**)). In addition to their role in autoinhibition, HR1, HR2, and HR3 interact with the fatty acyl chains of POPC in our simulations, suggesting that they cooperate in retaining lipid substrates near the catalytic center (Fig. 3(**e**–**g**)). Both HR2 and HR3 remained stably embedded in the membrane and acted as a firm scaffold from which a lipid acyl chain could be lifted toward the catalytic site, consistent with the complete loss of activity observed in the HR2 and HR3 mutants. By contrast, HR1 displayed greater flexibility, occasionally disengaging from the membrane and thereby contributing to conformational changes that promote opening of the catalytic site (Fig. 3(**a**–**c**)).

### Curvature dependence of Atg15 activity

Our findings reveal that Atg15 activity is influenced by membrane curvature (Fig. 5). Biochemical assays demonstrated that the cleavage of NBD-labeled phospholipids was markedly more efficient in 50–100 nm liposomes than in larger vesicles (Fig. 5(**a**)), indicating a preference for highly curved membranes. This curvature dependence is consistent with the results of our MD simulations using 50 nm POPC vesicles, which showed that Atg15 remained associated with positively curved membranes, whereas it tended to dissociate from negatively curved membranes and from flat bilayers (Fig. 5(**b**–**e**)). Together, these data indicate that Atg15 preferentially binds to positively curved membranes, where it more efficiently degrades membrane phospholipids. The curvature preference of Atg15 can be rationalized by differences in membrane affinity. The spatial arrangement of the hydrophobic regions identified in our mutational analysis, HR2 and HR3, appears more compatible with a positively curved bilayer, favoring stable membrane association under such conditions. Moreover, all-atom MD simulations revealed that Atg15 placed on a flat bilayer occasionally induced local positive curvature in the surrounding lipids (Supplementary Fig. S8), although Atg15 did not appear to show the pronounced curvature-inducing capacity characteristic of BAR-domain proteins^13^. These observations suggest that Atg15 preferentially binds to positively curved membranes and that membrane association stabilizes an open, catalytically competent state, thereby enhancing phospholipid hydrolysis. Accordingly, smaller intravacuolar vesicles such as Cvt vesicles (∼150 nm) and MVB intraluminal vesicles, which are smaller than Cvt vesicles, might represent more favorable substrates than the autophagic bodies (∼500 nm). Nevertheless, Atg15 efficiently disrupts Cvt vesicles, internal vesicles of the MVB, and autophagic bodies *in vivo*. In addition to positive curvature, other factors may also contribute to Atg15 activity and selectivity.

Curvature sensitivity may also contribute to protecting the vacuolar limiting membrane, which is nearly flat or even negatively curved and thus disfavored for stable Atg15 association (Fig. 5(**c**, **d**)). However, the vacuolar membrane can occasionally form inward invaginations or tubular structures that locally generate pronounced positive curvature. These regions would, in principle, be susceptible to Atg15. Yet the vacuolar membrane is rarely degraded, and our localization analyses do not detect Atg15 on the vacuolar limiting membrane^11^. Together, these observations suggest that curvature-dependent recognition contributes to Atg15 selectivity, but additional regulatory mechanisms that limit Atg15 access and/or activation at the vacuolar membrane are likely required to fully account for both its broad activity against intravacuolar vesicles and the protection of the vacuolar limiting membrane *in vivo*.

### Multilayer regulation of Atg15 activity ensures selective autophagic membrane degradation

Membrane degradation by autophagy is a universal process, yet it remains unclear whether a common molecular mechanism underlies this event across species. In yeast, the vacuole is much larger than the autophagosome, whereas in mammals autophagosomes are substantially larger than lysosomes. Although candidate lipases with phospholipase B activity have recently been identified as mediators of autophagosomal inner membrane degradation in mammals^14^, it is still unclear why lysosomal membranes are not degraded by such lipases, as is the case in yeast. Clarifying whether a shared principle, such as curvature-dependent recognition, governs autophagic membrane degradation across species will be an important subject for future investigation.

In this study, we elucidated how a potentially deleterious lipase, Atg15, exerts its activity by integrating MD simulations with biochemical experiments. Our findings not only provide insights into Atg15 function but also have broader implications for the study of phospholipases in general. Importantly, we show through MD simulations at the molecular level that Atg15 activity is controlled through multiple layers of regulation, including proteolytic processing within the vacuole and curvature-dependent interactions with membrane lipids, thereby ensuring that its lipolytic potential is unleashed only under appropriate conditions (Fig. 6). This work highlights the elaborate regulatory mechanisms that safeguard membrane degradation in cells and underscores the broader significance of Atg15 as a model for understanding membrane lipid degradation.

**Figure 6.**
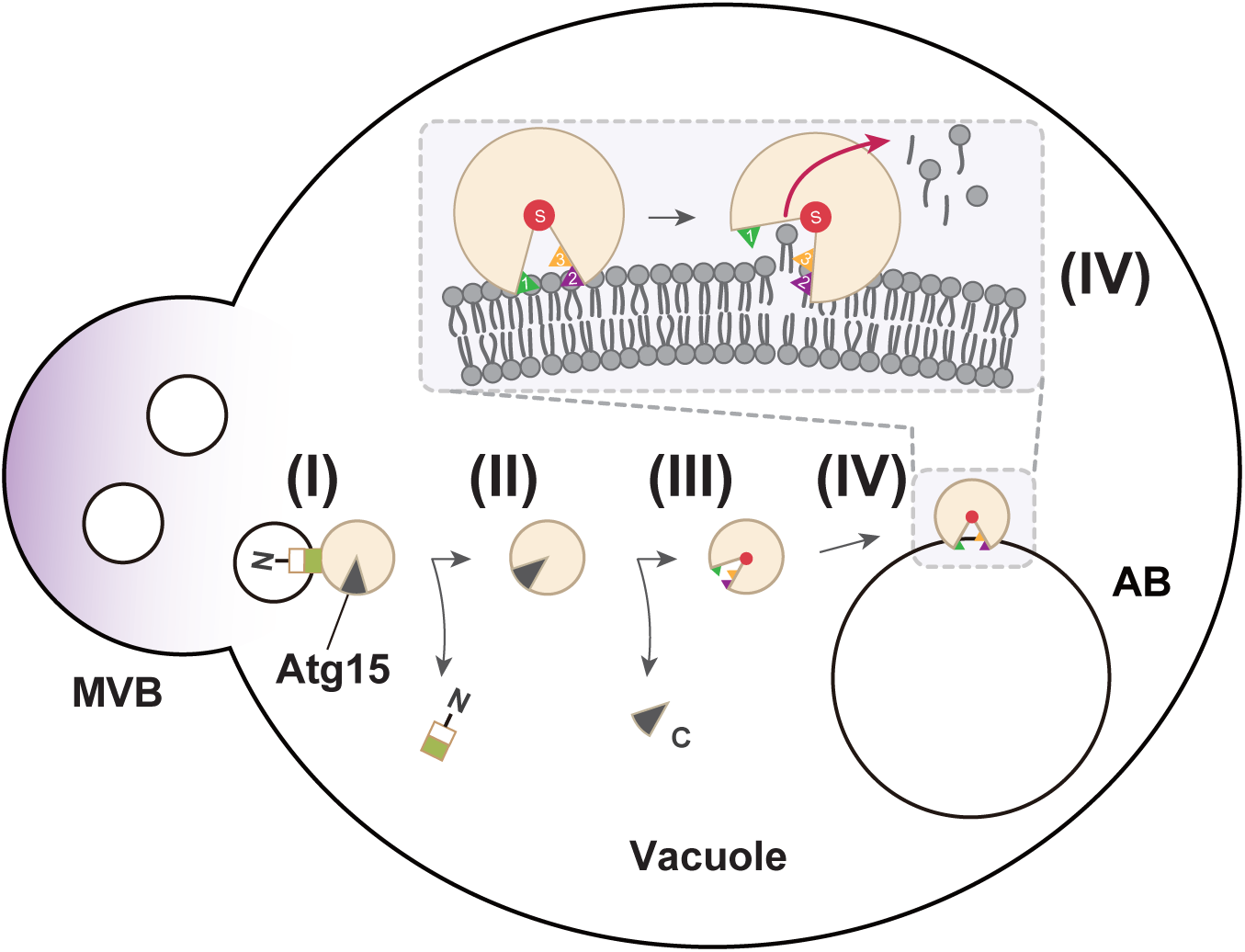
Proposed activation mechanism of Atg15 within the vacuole. Atg15 is delivered to the vacuole via the multivesicular body (MVB) pathway as the inactive precursor form I. Upon delivery, the N-terminal region of Atg15 is first cleaved by Pep4/Prb1, producing form II and releasing Atg15 from the MVB membrane. Subsequent cleavage at the C-terminal region generates form III, in which the catalytic pocket becomes partially exposed. When Atg15 associates with a lipid bilayer, its catalytic pocket opens further (form IV), allowing recognition and hydrolysis of lipid substrates. Three hydrophobic regions required for conformational opening, membrane association, and substrate recognition are indicated as HR1 (green), HR2 (purple), and HR3 (orange).

## Acknowledgements

We are grateful to members of the Ohsumi laboratory for discussions and critical comments, and the Integrative Bioscience Facility at Institute of Science Tokyo for DNA sequencing. This work was supported in part by JSPS KAKENHI Grant Number JP25K09559 (to T.K. and Y.S.), JP19H05707, JP23K20044, JP23K06667, JP24H00060, JP25H00966, JP25H01320, JP25H01321 (to N.N.N.), JP24K01980, JP23K05715 (to Y.S), Grant-in-Aid for Scientific Research on Innovative Areas JP22H04640 (T.K.), JP19H05708 (Y.O.), CREST, Japan Science and Technology Agency Grant number JPMJCR20E3 (to N.N.N.), and grants from the Takeda Science Foundation (to N.N.N.). T.K. was also supported by the JST FOREST program (JPMJFR214X). The numerical computations were performed with the supercomputer systems Fugaku (hp230252, hp250019) and HOKUSAI at RIKEN, Wisteria/BDEC-01 at the University of Tokyo.

## Author contributions

T.K., N.N.N, and Y.S. designed the research. N.N.N. and Y.S. performed structural analysis. Y.S. performed MD simulations. T.K. and M.S. performed experiments. T.K., N.N.N, and Y.S. analyzed data. T.K., N.N.N., Y.O., and Y.S. wrote the manuscript with the inputs from all the authors. T.K. and Y.S. supervised the work.

## Declaration of interests

The authors declare no competing interests.

## Materials and Methods

### Structural prediction using AlphaFold2

The structure of Atg15 in Fig. 1(**a**) was predicted using AlphaFold2 v2.3 installed on a local computer (Sunway Technology Co., Ltd., Tokyo, Japan)^15^. The predictions were run with five models and a single seed per model, and default multiple sequence alignment generation using the MMSeqs2 server^16^. The unrelaxed predicted models were subjected to an Amber relaxation procedure and the relaxed model with the highest confidence based on predicted LDDT scores was selected as the best model and used for MD simulation and figure preparation^15^. Structural figures were prepared using PyMOL (http://www.pymol.org/pymol)^17^.

### Lipids used in this study

NBD-PE (1-oleoyl-2-[12-[(7-nitro-2-1,3-benzoxadiazol-4-yl)amino]dodecanoyl]-*sn*-glycero-3-phosphoethanolamine), NBD-PC (1-Oleoyl-2-[12-[(7-nitro-2-1,3-benzoxadiazol-4-yl)amino]dodecanoyl]-sn-Glycero-3-Phosphocholine), DOPE (dioleoylphosphatidylethanolamine), POPC (1-palmitoyl-2-oleoylphosphatidylcholine) were obtained from Avanti Polar Lipids. For lipase assays performed in solution, NBD-PE was dissolved in 5 mM CHAPS at a concentration of 125 ng/µL and sonicated for 5 min.

### Yeast Strains and media

The yeast strains used in this study are listed in Table S1. Gene deletion and tagging were performed using standard PCR-based methods, as described previously^18,19^, and validated by PCR. Cells were cultured at 30°C in YPD medium (1% yeast extract (Gibco), 2% Bacto peptone (Gibco), 2% glucose) or in a synthetic defined medium comprising 0.17% yeast nitrogen base without amino acids and ammonium sulfate (BD), 0.5% ammonium sulfate, 0.5% casamino acids (BD), 2% glucose, 20 µg/mL adenine sulfate, and 20 µg/mL tryptophan (SD/CA medium). To induce autophagy, cells were treated with rapamycin (LC Laboratories, R-5000) at a concentration of 0.2 µM or cultured in SD-N medium.

### Plasmid construction

Plasmids used in this study are listed in Table S2. Plasmids *pRS426-ATG15(Y276D, L277D, W278D, 168-3xFLAG-169)* (SMP81), *pRS426-ATG15(L403D, V404D, G405D, Y406D, 168-3xFLAG-169)* (SMP149), and *pRS426-ATG15(G246D, L247D, P248D, G249D, 168-3xFLAG-169)* (SMP116) were generated from *pRS426-ATG15(168-3xFLAG-169)* (YPL183) by site-directed mutagenesis using inverse PCR with PrimeSTAR Max (Takara) or KOD One (Toyobo). The plasmid *pRS426-CPY(1–50)-ATG15(51–471)* (SMP150) was constructed using *pRS426-CPY(1–50)-ATG15(36–479)* (YPL112) as a template. A DNA fragment encoding amino acids 51–471 of *ATG15* was amplified by PCR and inserted into a linearized vector lacking residues 36–50 and 472–479, which was amplified by PCR from the same template. The two fragments with overlapping ends were assembled using the In-Fusion cloning system (Clontech).

### Preparation of purified Atg15 for lipase activity assay

Purified Atg15 protein was obtained essentially as described previously^11^. Briefly, yeast cells overexpressing internally FLAG-tagged Atg15 were treated with rapamycin for 1 h, and vacuoles were subsequently isolated. The vacuolar pellet fraction was collected by centrifugation, solubilized in buffer containing 0.5% n-dodecyl-β-D-maltoside (DDM), and FLAG-tagged Atg15 was affinity-purified using anti-FLAG agarose beads. Bound proteins were competitively eluted with 3×FLAG peptide, and the eluates were directly subjected to the lipase assay.

### Liposome preparation

For liposome-based assays, lipid mixtures of either POPC:NBD-PC (94:6, v/v) or POPC:DOPE:NBD-PE (88:6:6, v/v) that were dissolved in chloroform and transferred to glass tubes. Dried lipid films were obtained by evaporating chloroform. The films were hydrated in buffer E (30 mM MES-Tris, pH 6.9, 200 mM KCl) to yield 1 mM lipid suspensions, which were incubated at room temperature for 1 h with intermittent vortexing. Homogeneously sized unilamellar vesicles were generated by sequential extrusion through polycarbonate membranes with pore sizes of 1000 (code no. 610010 Avanti), 400 (code no. 610007 Avanti), 100 (code no. 610005 Avanti), and 50 nm (code no. 610003 Avanti) using a mini-extruder (Avanti). To remove free (non-incorporated) fluorescent lipids, liposome suspensions containing NBD-PC or NBD-PE were centrifuged at 125,000 × g for 1 h at 4°C, and the supernatants were carefully discarded. The pellets were resuspended in buffer E, and liposome size distribution was confirmed using a Zetasizer Nano S (Malvern Instruments)^8^.

### Lipase activity assays

Lipase activity assays of purified Atg15 were performed as follows. For assays testing the effect of DTT (Fig. 1(**c**)), purified Atg15 was incubated in buffer E (20 μL) with or without 5 mM DTT at room temperature, 40°C, or 50°C for 5 min. After this pre-incubation, the samples were placed on ice to cool, and then 125 ng of free NBD-PE was added to initiate the reaction. Reactions were carried out at 30°C for 2 h. For assays using liposomes, 1 μL of liposomes (1 mM) containing NBD-PE or NBD-PC was added to 20 μL buffer E, and then mixed with purified Atg15 for 2 h at 30°C. After incubation, total lipids were extracted using the BUME method^11^, dissolved in chloroform/methanol (2:1, v/v), and applied to HPTLC silica gel 60 F254 plates (Millipore, 1.05642.0001). Plates were developed in chloroform/methanol/water (65:35:8, v/v/v) as described previously^20^. Fluorescence was detected with a FUSION-FX7 imaging system (Vilber-Lourmat), and band intensities were quantified using the corresponding analysis software.

### Western blotting

Immunoblot analyses were performed essentially as described previously^8,11^. The following primary antibodies were used: α-FLAG (Sigma–Aldrich, F3165, 1:1000), α-Pho8 (Abcam, ab113688, 1:1000), α-Ape1 (laboratory stock, 1:5000), α-Prb1 (laboratory stock, 1:5000), α-Atg15 (laboratory stock, 1:1000), and α-GFP (Roche, 11814460001, 1:1000). Chemiluminescence was detected with Femtoglow HRP Substrate (Michigan Diagnostics) and visualized using a FUSION-FX7 imaging system.

### MD simulation

Simulations of Atg15 and its processed forms in solution and on flat membranes were performed using GENESIS^21^ and the CHARMM36 force field^22^. The initial structures of the proteins were predicted using AlphaFold2. For simulations of the proteins in solution, the proteins were solvated in a 10 nm × 10 nm × 10 nm box containing TIP3P water molecules and 0.15 M KCl ions. For the calculation of the proteins on the flat membrane, the membranes consisting of 300 POPC lipid molecules in each leaflet were used. The protein and lipids were solvated in a 15 nm×15 nm×14 nm box with TIP3P waters and 0.15 M KCl ions. The locations of Atg15 and its processed forms on the membranes were predicted using the PPM server^12^. The initial configurations were built by the Membrane Builder module in CHARMM-GUI server^23^. Energy minimization was performed for 1000 steps by the steepest descent algorithm and then by the conjugate gradient algorithm. Then a 250 ps NVT simulation was performed at 323.15 K for solvent equilibration, followed by a 1.6 ns NPT equilibration to 1 atm using the Langevin thermostat/barostat^24^. The production MD simulations were performed for 100 ns with a time-step of 2 fs and Langevin thermostat/barostat. Long-range electrostatic interactions were treated by the particle-mesh Ewald method^25,26^. The short-range electrostatic and van der Waals interactions both used a cutoff of 12 Å. All bonds were constrained by the SHAKE/RATTLE algorithm^27,28^.

Simulations of Atg15 and its processed forms on POPC vesicles of 50 nm diameter were performed using a coarse-grained model with GROMACS (version 2022)^29^ and the MARTINI model (version 2.2)^30,31^. The location of the proteins on the membranes was predicted using the PPM server^12^. The tertiary structure of the protein in the coarse-grained model was maintained using an elastic network model, employing the open-state structure obtained from a 100 ns simulation of Atg15 Form III with disulfide bonds on the membrane using an all-atom model. The proteins and lipids were solvated in a 65 nm × 65 nm × 65 nm box with coarse-grained waters and 0.15 M NaCl ions. The initial configurations were built by the Martini Maker module in CHARMM-GUI server^23^. Each system was first energy minimized and equilibrated using the Berendsen thermostat and barostat^32^ followed by 1 μs production runs with a 20-fs time step along with position restraints on the protein. System temperature and pressure during the production phase were maintained at 323.15 K and 1 atm with the velocity rescaling thermostat and the semi-isotropic Parrinello–Rahman barostat^33^, respectively. The simulation results were visualized and analyze using PyMOL (http://www.pymol.org/pymol).

## Supplementary Figure legends

**Supplementary Figure S1. Detailed view of the D51 and S471 regions in Atg15 Form I predicted by AlphaFold.**

Atg15 is represented in a wheat-colored cartoon, with the D51 and S471 areas highlighted in blue.

**Supplementary Figure S2. All-atom MD simulation of Atg15 (Form III with disulfide bonds) in solution**.

Five additional simulation results independently calculated from the same initial state as Fig. 2(**a**). The three left panels represent the structures at 10, 50 (or 80) and 100 ns, respectively. Each representation is the same as in Fig. 2(**a**). The right panel shows the time dependence of the distance D_13_.

**Supplementary Figure S3. All-atom MD simulation of Atg15 (Form II with disulfide bonds) in solution. (a–d)**.

Four additional simulation results independently calculated from the same initial state as Fig. 2(**c**). The three left panels represent the structures at 10, 50 and 100 ns, respectively. Each representation is the same as in Fig. 2(**c**). The right panel shows the time dependence of the distances D_13_ and D_14_.

**Supplementary Figure S4. Structural changes of Atg15 (form II with disulfide bonds) in solution**.

(**a**) Overall views of Atg15 at 0 and 100 ns, respectively. (**b**) Detailed views around residue S332 at 0 and 100 ns, respectively. Each representation is the same as in Fig. 2(**c**).

**Supplementary Figure S5. All-atom MD simulation of Atg15 (Form III without disulfide bonds) in solution**.

Four additional simulation results independently calculated from the same initial state as Fig. 2(**e**). The three left panels represent the structures at 10, 50 and 100 ns, respectively. Each representation is the same as in Fig. 2(**e**). The right panel shows the time dependence of the distance D_13_.

**Supplementary Figure S6. All-atom MD simulation of structural changes in Atg15 (Form III with disulfide bonds) on a POPC bilayer**.

Four additional simulation results independently calculated from the same initial state as Fig. 3(**a**). The three left panels represent the structures at 10, 50 (or 75) and 100 ns, respectively. Each representation is the same as in Fig. 3(**a**) top. The right panel shows the time dependence of the distance D_13_.

**Supplementary Figure S7. All-atom MD simulation of membrane interaction in Atg15 (Form III with disulfide bonds) on a POPC bilayer**.

Four additional simulation results independently calculated from the same initial state as Fig. 3(**a**). The three left panels represent the structures at 10, 50 (or 75) and 100 ns, respectively. Each representation is the same as in Fig. 3(**a**) bottom. The right panel shows the time dependence of the distance D_lip_.

**Supplementary Figure S8. Curvature changes of POPC membranes associated with Atg15 (form III with disulfide bonds)**.

Snapshots at 70, 75, and 80 ns, respectively, from the same simulation as Fig. 3(**a**). Phosphorus and the others of POPC bilayer are represented as orange spheres and green lines, respectively. Other representations are the same as in Fig. 3(**a**) bottom.

**Supplementary Figure S9. All-atom MD simulation of Atg15 (Form III with disulfide bonds) with a POPC molecule in solution**.

Four additional simulation results independently calculated from the same initial state as Fig. 3(**e**). The two left panels represent the structures at 10 and 100 ns, respectively. Each representation is the same as in Fig. 3(**e**). The two right panels show the time dependence of the distances D_13_ and D_sn_, respectively.

**Supplementary Figure S10. All-atom MD simulation of Atg15 HR1D mutant in solution**.

Four additional simulation results independently calculated from the same initial state as Fig. 4(**b**). The three left panels represent the structures at 10, 50 and 100 ns, respectively. Each representation is the same as in Fig. 4(**b**). The right panel shows the time dependence of the distance D_13_.

**Supplementary Figure S11. All-atom MD simulation of Atg15 HR2D mutant in solution**.

Four additional simulation results independently calculated from the same initial state as Fig. 4(**d**). The three left panels represent the structures at 10, 50 and 100 ns, respectively. Each representation is the same as in Fig. 4(**d**). The right panel shows the time dependence of the distance D_13_.

**Supplementary Figure S12. All-atom MD simulation of Atg15 HR3D mutant in solution**.

Four additional simulation results independently calculated from the same initial state as Fig. 4(**f**). The three left panels represent the structures at 10, 50 and 100 ns, respectively. Each representation is the same as in Fig. 4(**f**). The right panel shows the time dependence of the distance D_13_.

**Supplementary Figure S13. Evaluation of Atg15 lipase activity using liposomes of various sizes**.

**(a)** Dynamic light scattering (DLS) profiles of the liposomes used for the lipase assay in Fig. 5(**a**). **(b)** DLS profiles of the liposomes used for the lipase assay in Supplementary Fig. S13(**c**). **(c)** Lipase activity of purified Atg15 toward liposomes containing NBD-PE. POPC liposomes containing 6% DOPE and 6% NBD-PE were prepared by sequential extrusion through polycarbonate filters with nominal pore sizes of 1000, 400, and 100 nm. The liposomes were incubated with purified wild-type Atg15 or Atg15(S332A), and the reactions were analyzed by TLC. The amounts of NBD-PE, NBD-LPE, and NBD-FFA were quantified. The relative proportions of each lipid species were calculated, and the differences between wild-type Atg15 and Atg15(S332A) were expressed as ΔRelative intensity.

**Supplementary Figure S14. Coarse-grained MD simulations of the Atg15 Form III on membranes with different curvatures**.

Time dependence of distance D_s_ for four additional independent calculations of Atg15 initially positioned on the positively curved outer surface (**a**), on the negatively curved inner surface (**b**) of a 50 nm diameter POPC vesicle and positioned on a flat POPC bilayer (**c**). The initial structures of panels (**a**–**c**) are the same as those in Fig. 5(**b**–**d**), respectively.

**Supplementary Movie S1. All-atom MD simulations of Atg15 (Form III with disulfide bonds) in solution.** Each representation is the same as in Fig. 2(**a**).

**Supplementary Movie S2. All-atom MD simulations of Atg15 (Form II with disulfide bonds) in solution.** Each representation is the same as in Fig. 2(**c**).

**Supplementary Movie S3. All-atom MD simulations of Atg15 (Form III without disulfide bonds) with a POPC molecule in solution.** Each representation is the same as in Fig. 2(**e**).

**Supplementary Movie S4. All-atom MD simulations of Atg15 (Form III with disulfide bonds) on a POPC bilayer.** Each representation is the same as in Fig. 3(**a**) bottom.

**Supplementary Movie S5. All-atom MD simulations of Atg15 (Form III with disulfide bonds) with a single POPC molecule in solution.** Each representation is the same as in Fig. 3(**e**).

## Notes

### Competing Interest Statement

The authors have declared no competing interest.

