## Supplementary Figures for "Structural and mechanistic basis for membrane recognition and activation of the vacuolar lipase Atg15"

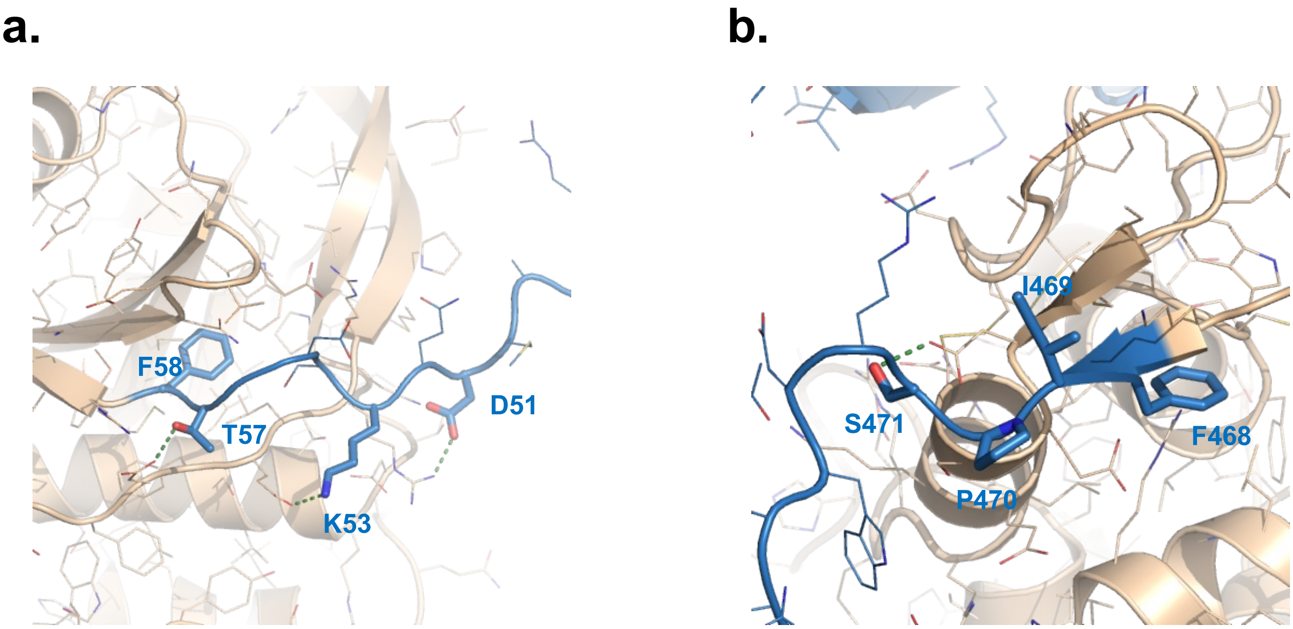


Figure. S1

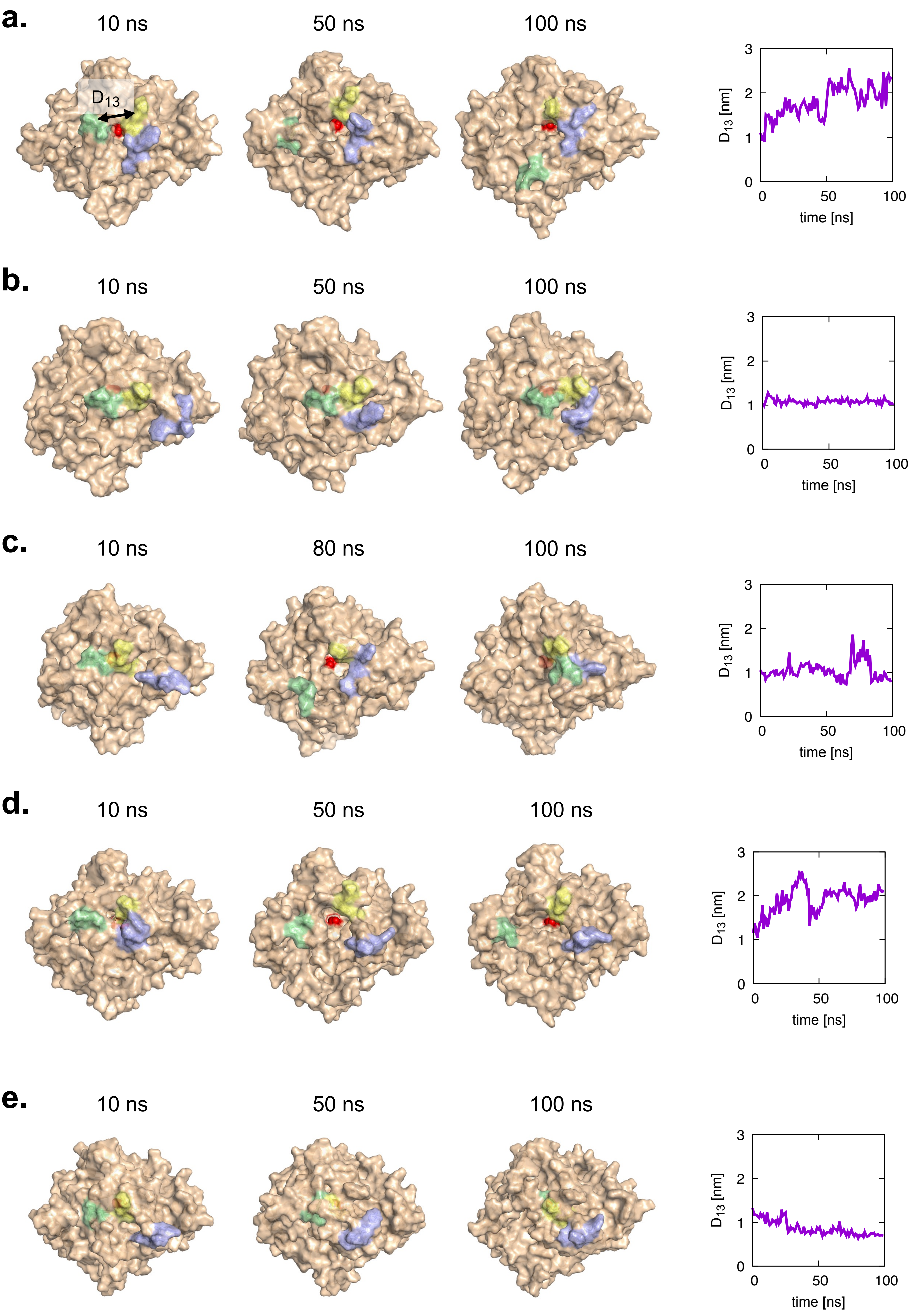

a.
10 ns
50 ns
100 ns
D13
D13 [nm]
time [ns]
b.
10 ns
50 ns
100 ns
D13 [nm]
time [ns]
c.
10 ns
80 ns
100 ns
D13 [nm]
time [ns]
d.
10 ns
50 ns
100 ns
D13 [nm]
time [ns]
e.
10 ns
50 ns
100 ns
D13 [nm]
time [ns]


Figure. S2

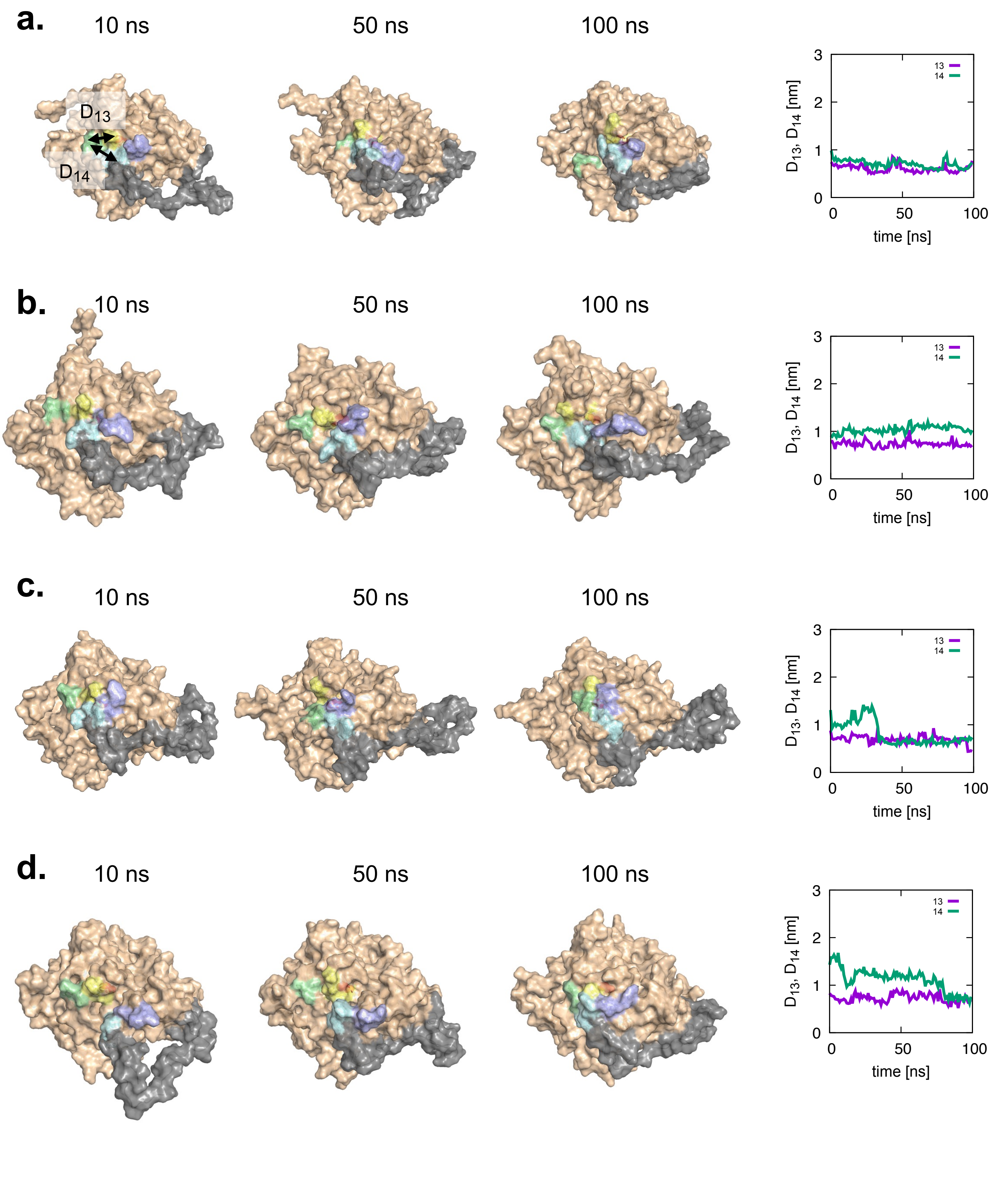

a.
10 ns
50 ns
100 ns
D13
D14
b.
10 ns
50 ns
100 ns
c.
10 ns
50 ns
100 ns
d.
10 ns
50 ns
100 ns
D13, D14 [nm]
time [ns]
13
14


Figure. S3

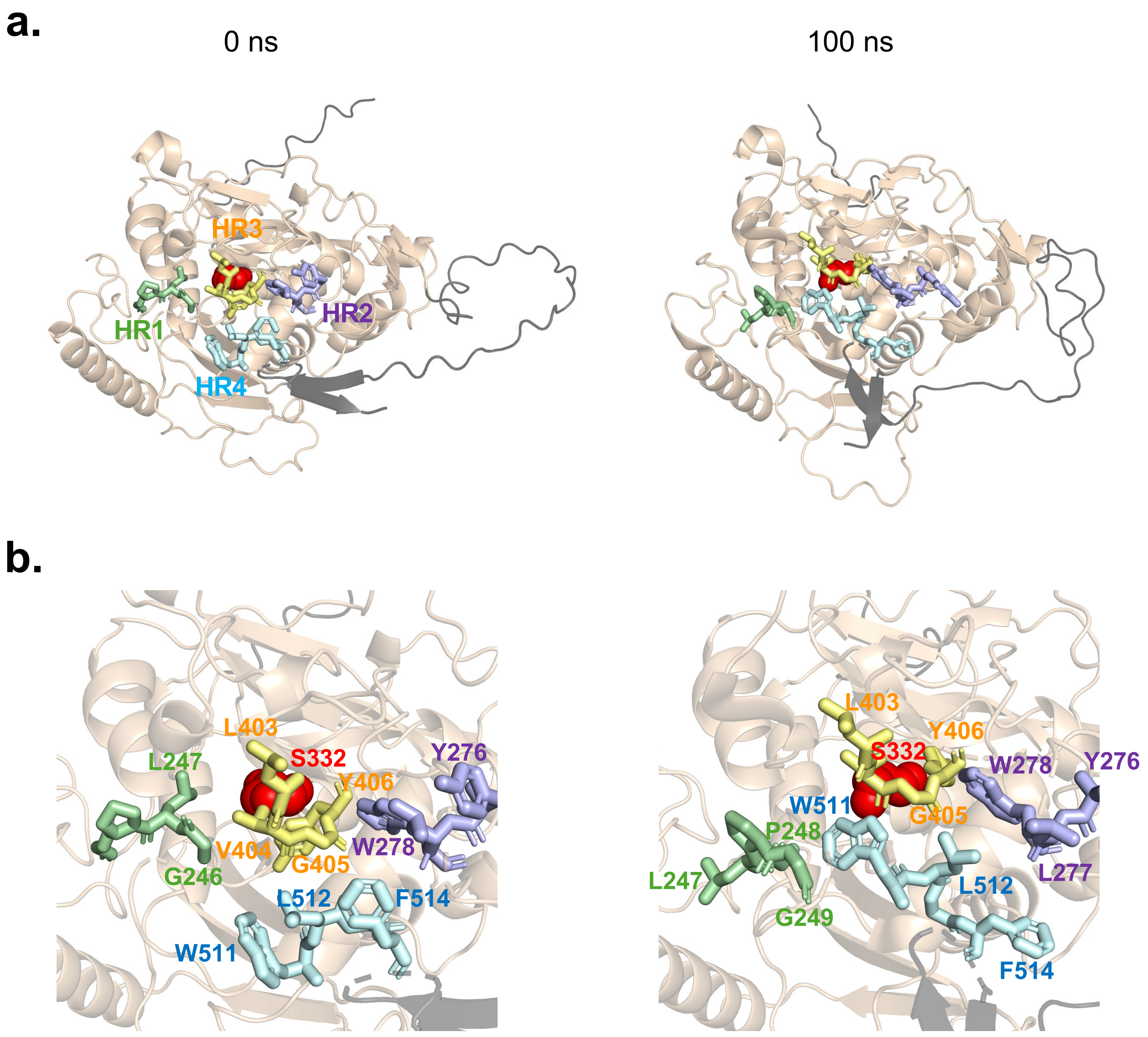
a.
0 ns
100 ns
HR3
HR1
HR2
HR4
b.
L403
S332
L247
Y276
Y406
V404
G405
W278
G246
L512
F514
W511
L403
Y406
S332
W278
Y276
W511
G405
P248
L247
L512
L277
G249
F514

Figure. S4

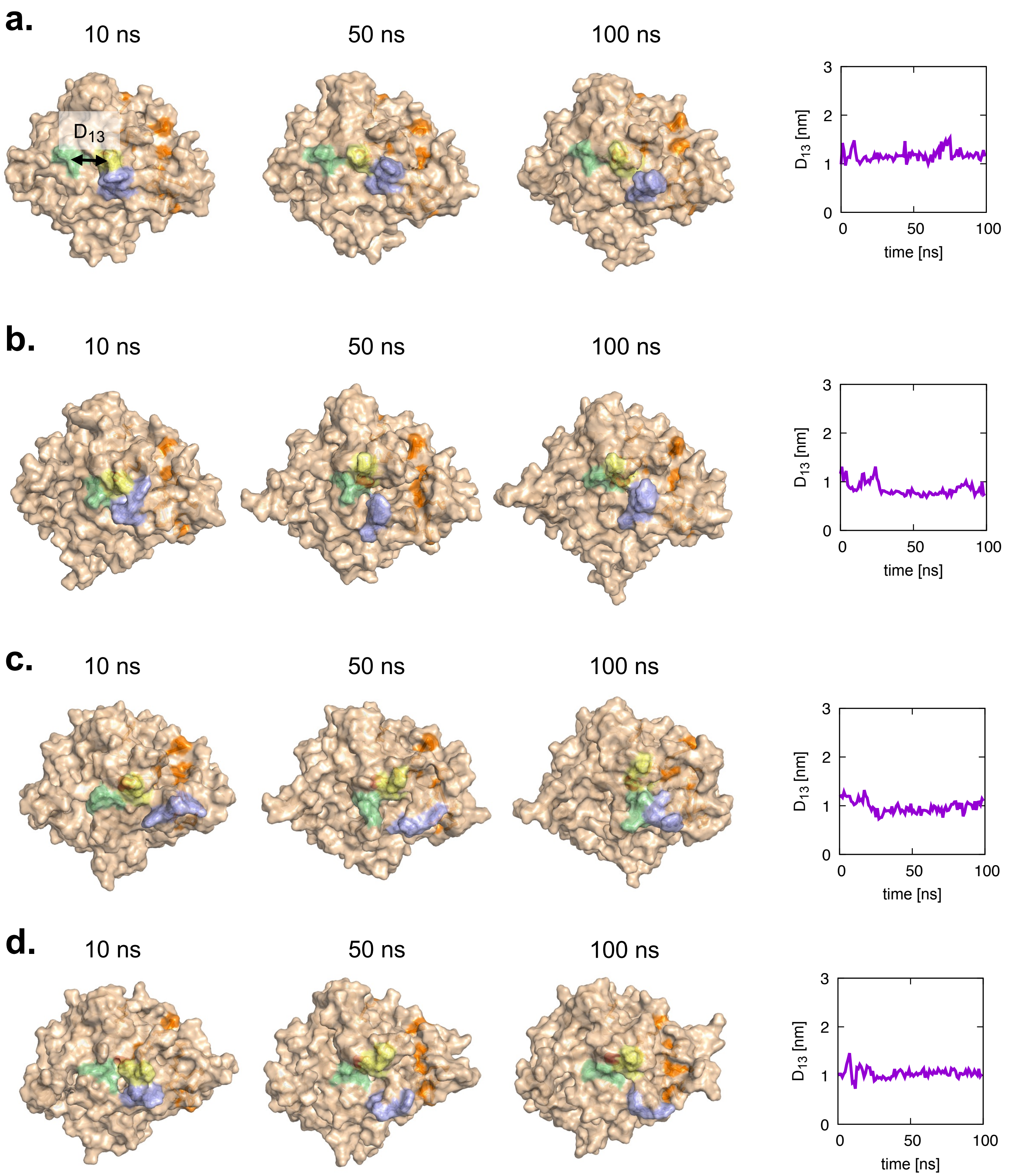

a.
10 ns
50 ns
100 ns
D13
b.
10 ns
50 ns
100 ns
c.
10 ns
50 ns
100 ns
d.
10 ns
50 ns
100 ns
D13 [nm]
time [ns]
0
1
2
3
0
50
100


Figure. S5

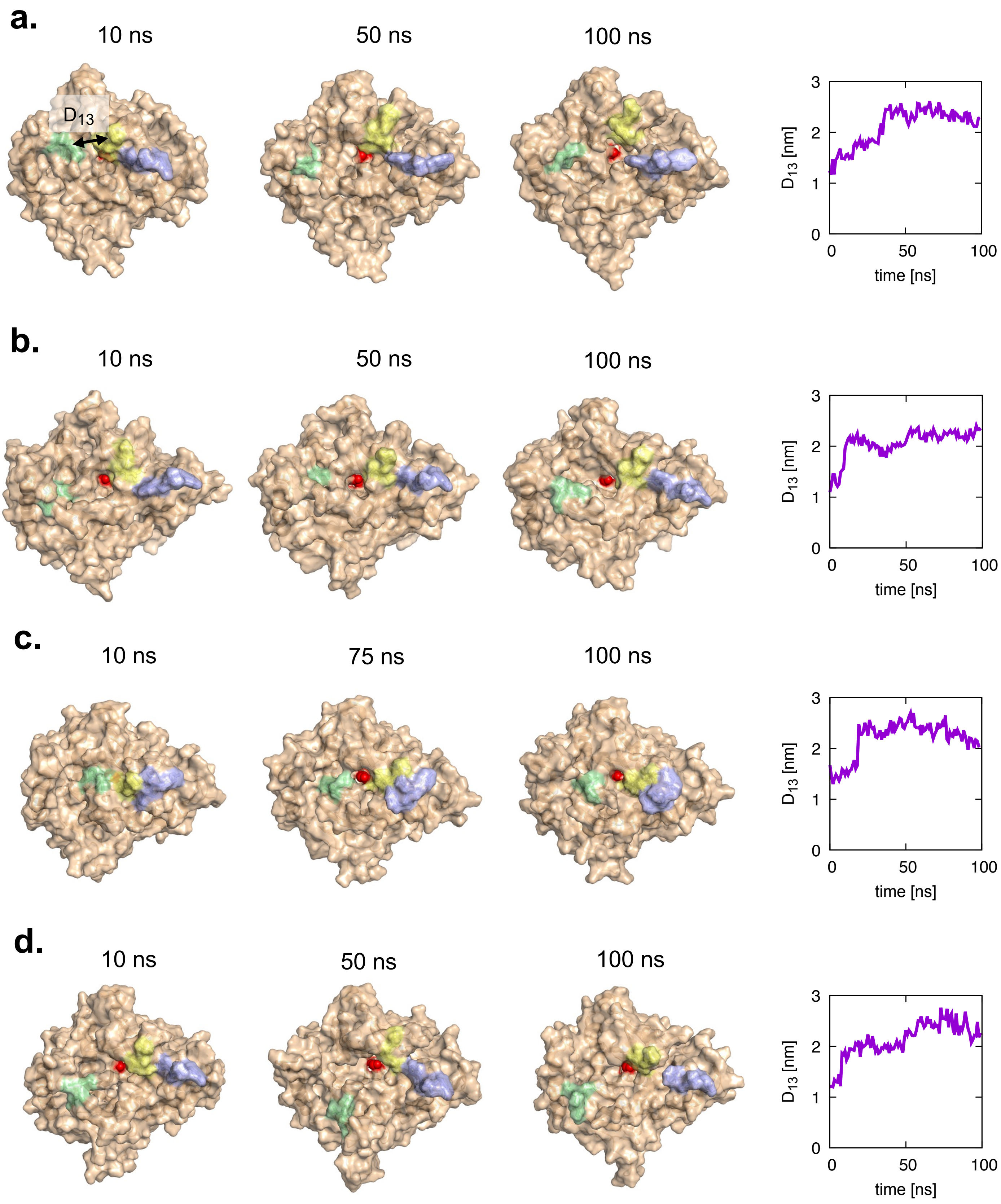
a.
10 ns
50 ns
100 ns
$D_{13}$
$D_{13}$ [nm]
time [ns]
b.
10 ns
50 ns
100 ns
$D_{13}$ [nm]
time [ns]
c.
10 ns
75 ns
100 ns
$D_{13}$ [nm]
time [ns]
d.
10 ns
50 ns
100 ns
$D_{13}$ [nm]
time [ns]

Figure. S6

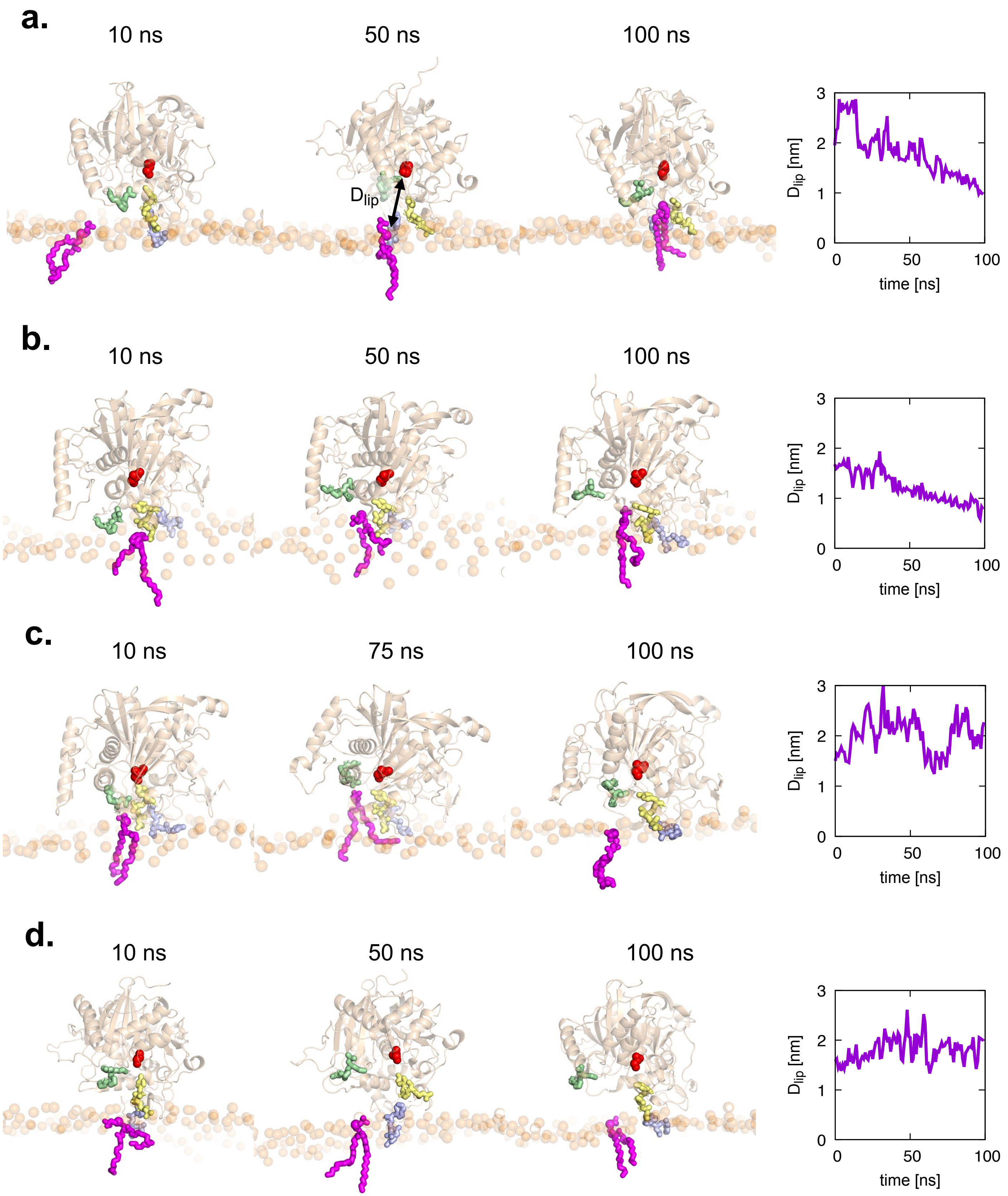

a.
10 ns
50 ns
100 ns
D$_{lip}$
D$_{lip}$ [nm]
time [ns]
0
50
100
0
1
2
3
b.
10 ns
50 ns
100 ns
D$_{lip}$ [nm]
time [ns]
c.
10 ns
75 ns
100 ns
D$_{lip}$ [nm]
time [ns]
d.
10 ns
50 ns
100 ns
D$_{lip}$ [nm]
time [ns]


Figure. S7

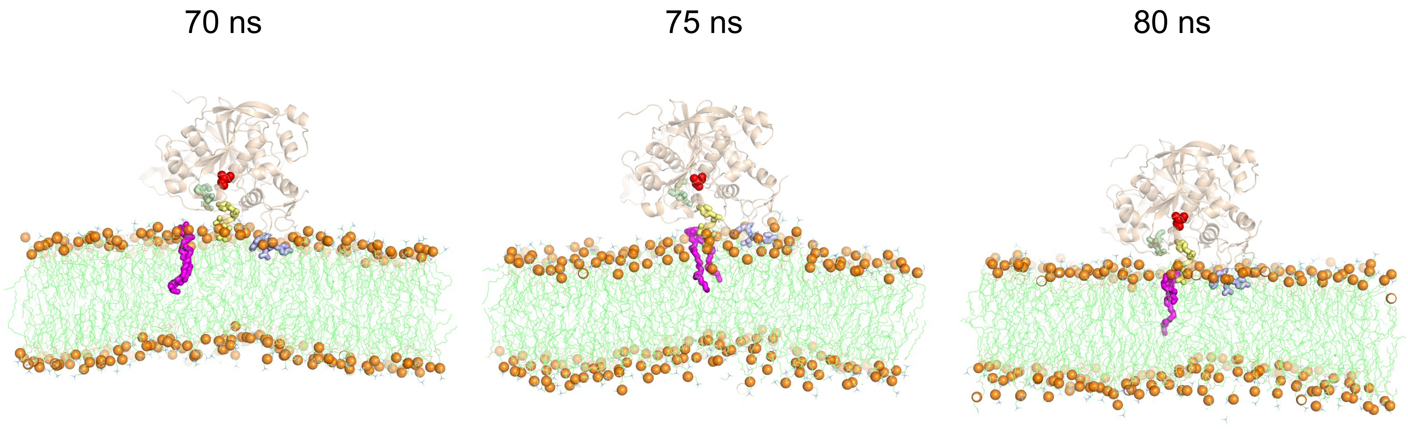


Figure. S8

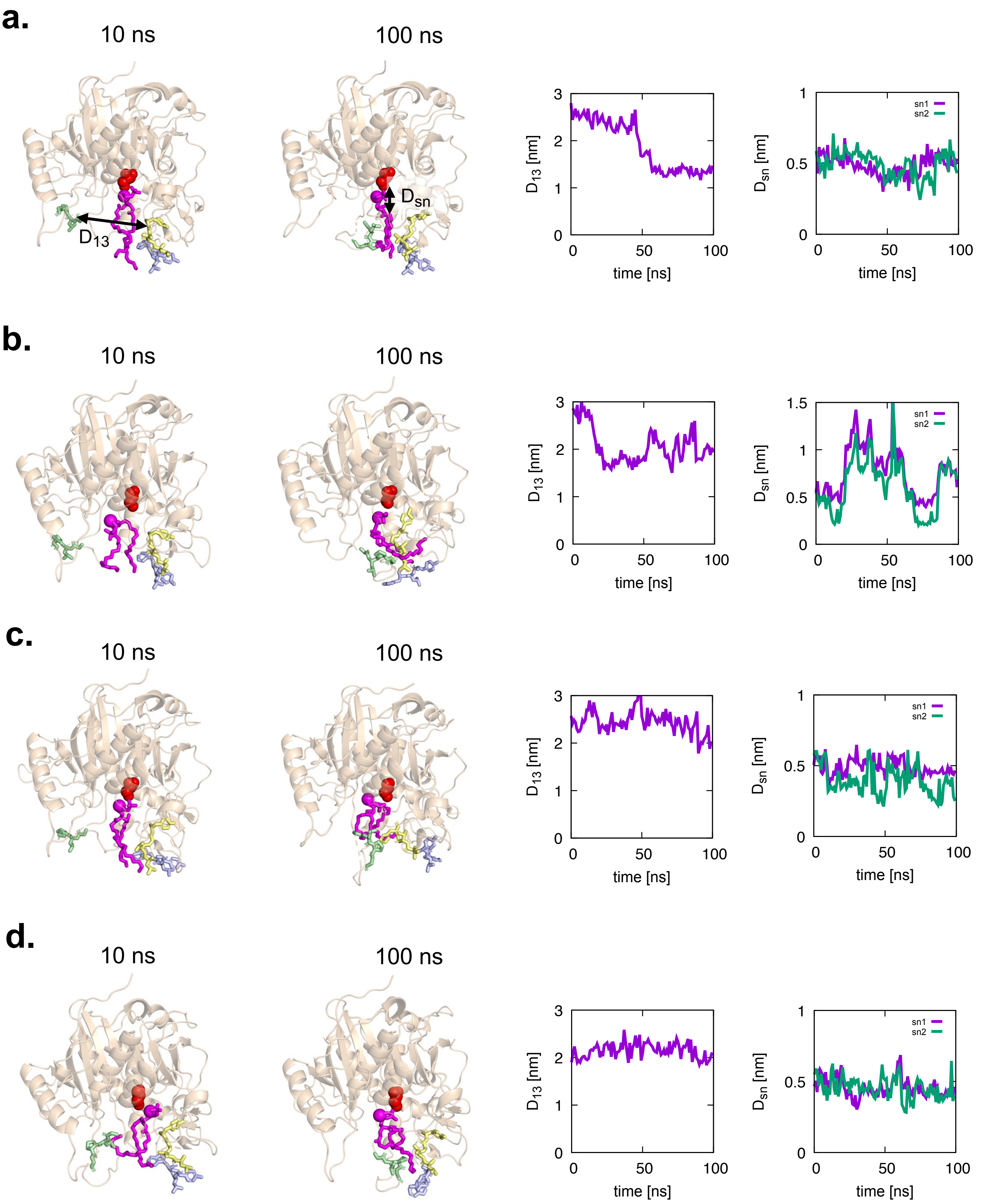
a.
10 ns
100 ns
D13
Dsn
D13 [nm]
time [ns]
Dsn [nm]
sn1
sn2
b.
10 ns
100 ns
c.
10 ns
100 ns
d.
10 ns
100 ns

Figure. S9

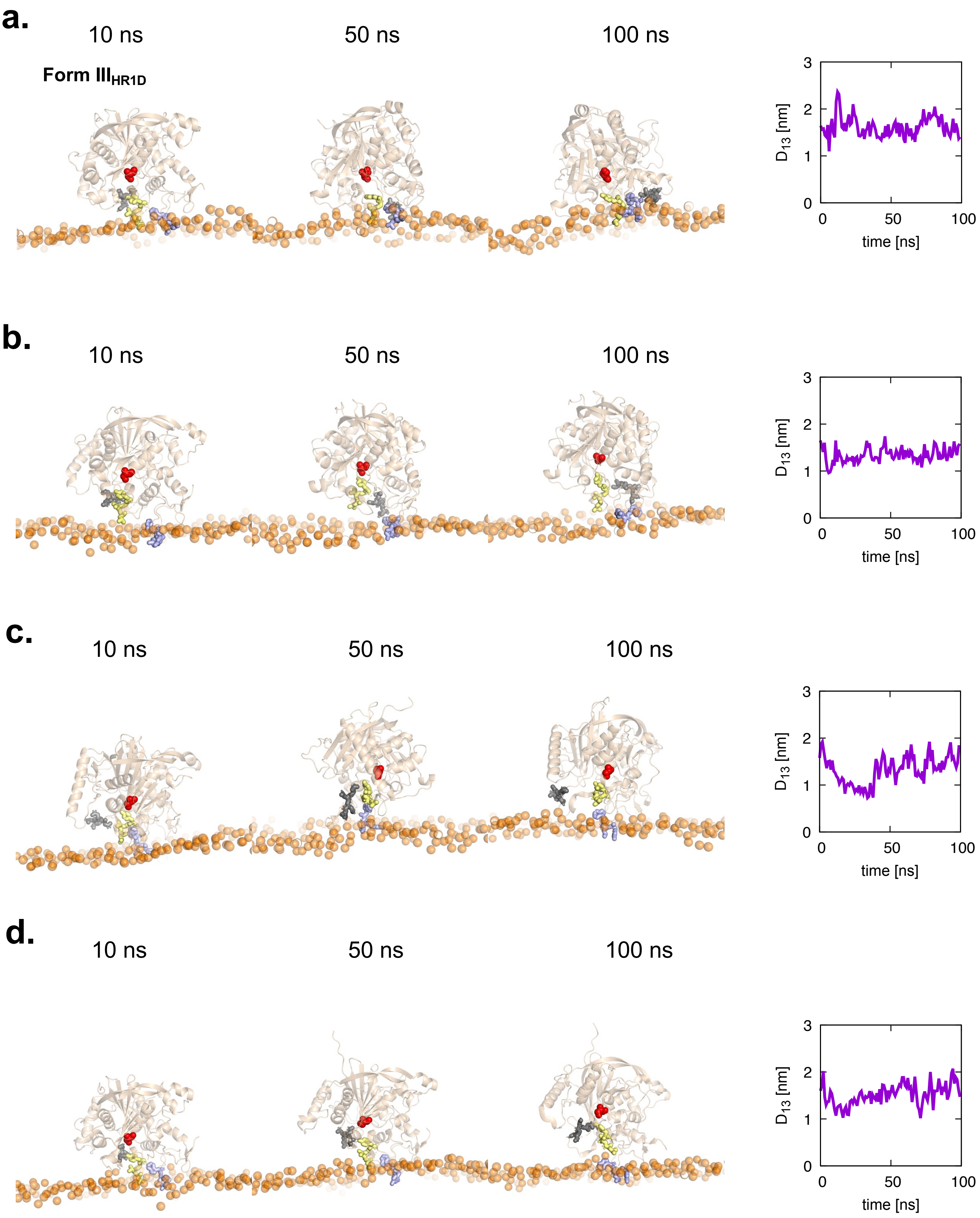
a.
10 ns
50 ns
100 ns
Form III$_{HR1D}$
$D_{13}$ [nm]
time [ns]
b.
10 ns
50 ns
100 ns
$D_{13}$ [nm]
time [ns]
c.
10 ns
50 ns
100 ns
$D_{13}$ [nm]
time [ns]
d.
10 ns
50 ns
100 ns
$D_{13}$ [nm]
time [ns]

Figure. S10

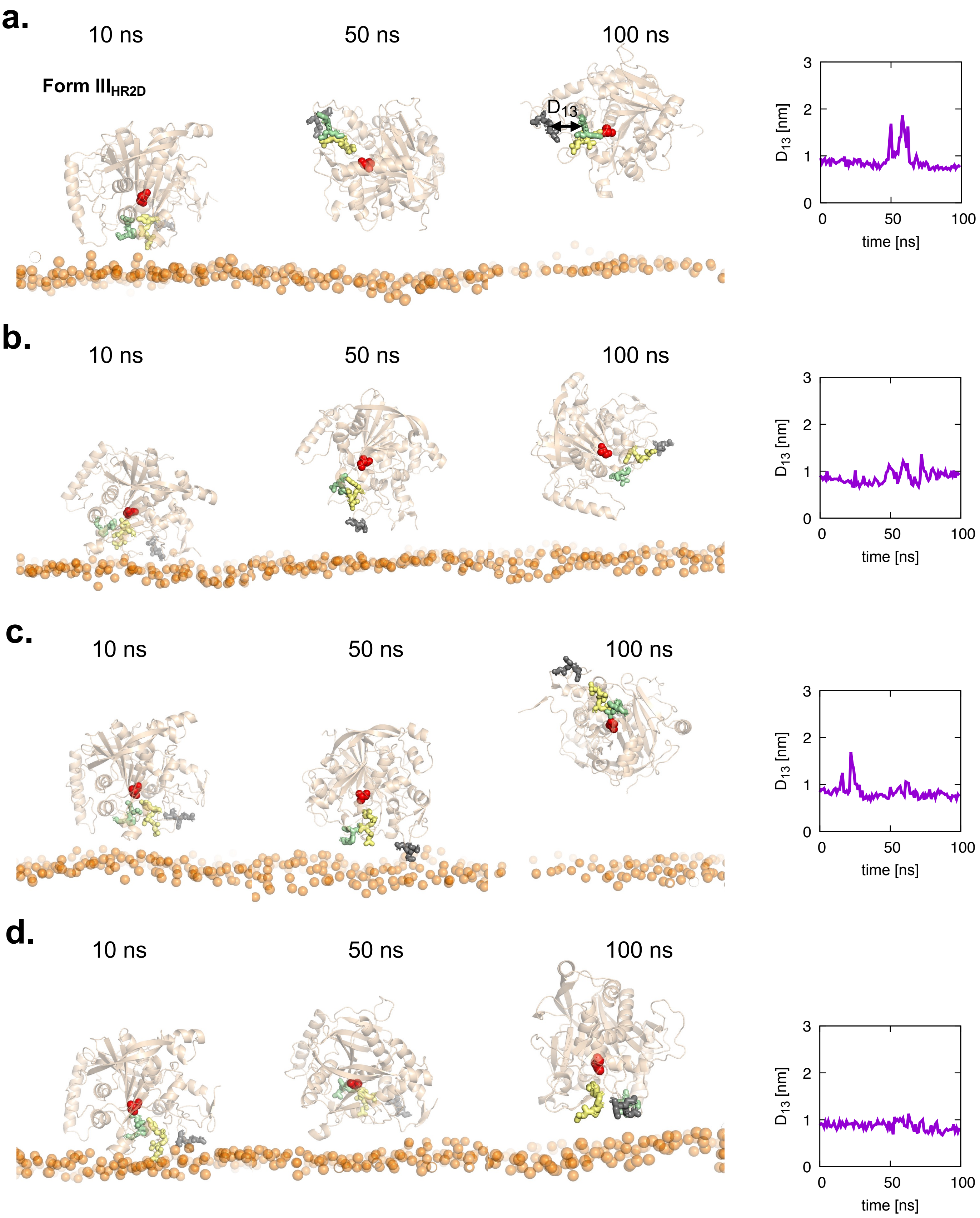
a.
10 ns
50 ns
100 ns
Form III$_{HR2D}$
$D_{13}$
$D_{13}$ [nm]
time [ns]
b.
10 ns
50 ns
100 ns
$D_{13}$ [nm]
time [ns]
c.
10 ns
50 ns
100 ns
$D_{13}$ [nm]
time [ns]
d.
10 ns
50 ns
100 ns
$D_{13}$ [nm]
time [ns]

Figure. S11

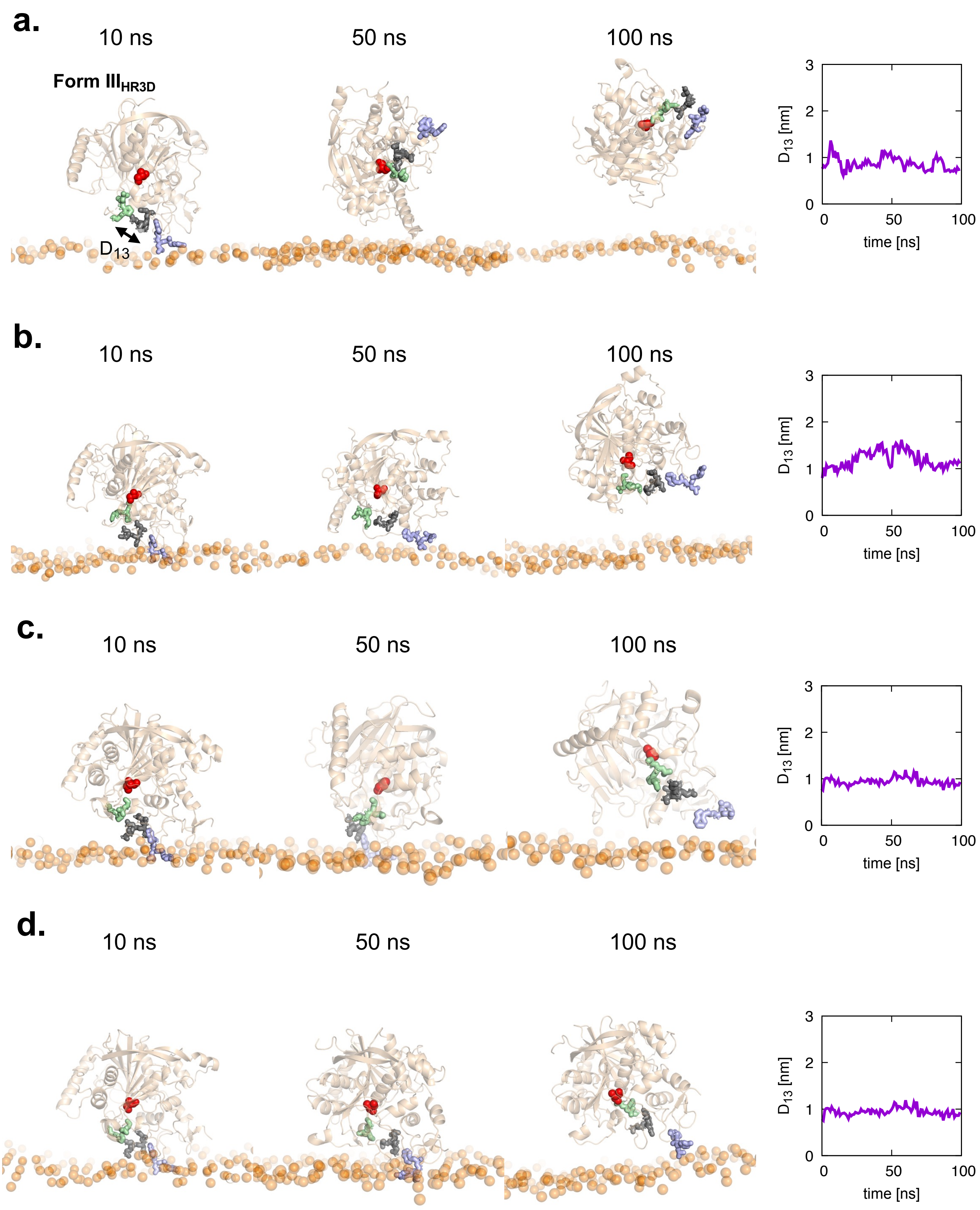
a.
10 ns
50 ns
100 ns
Form III$_{HR3D}$
D$_{13}$
b.
c.
d.
D$_{13}$ [nm]
time [ns]
0
1
2
3
50
100

Figure. S12

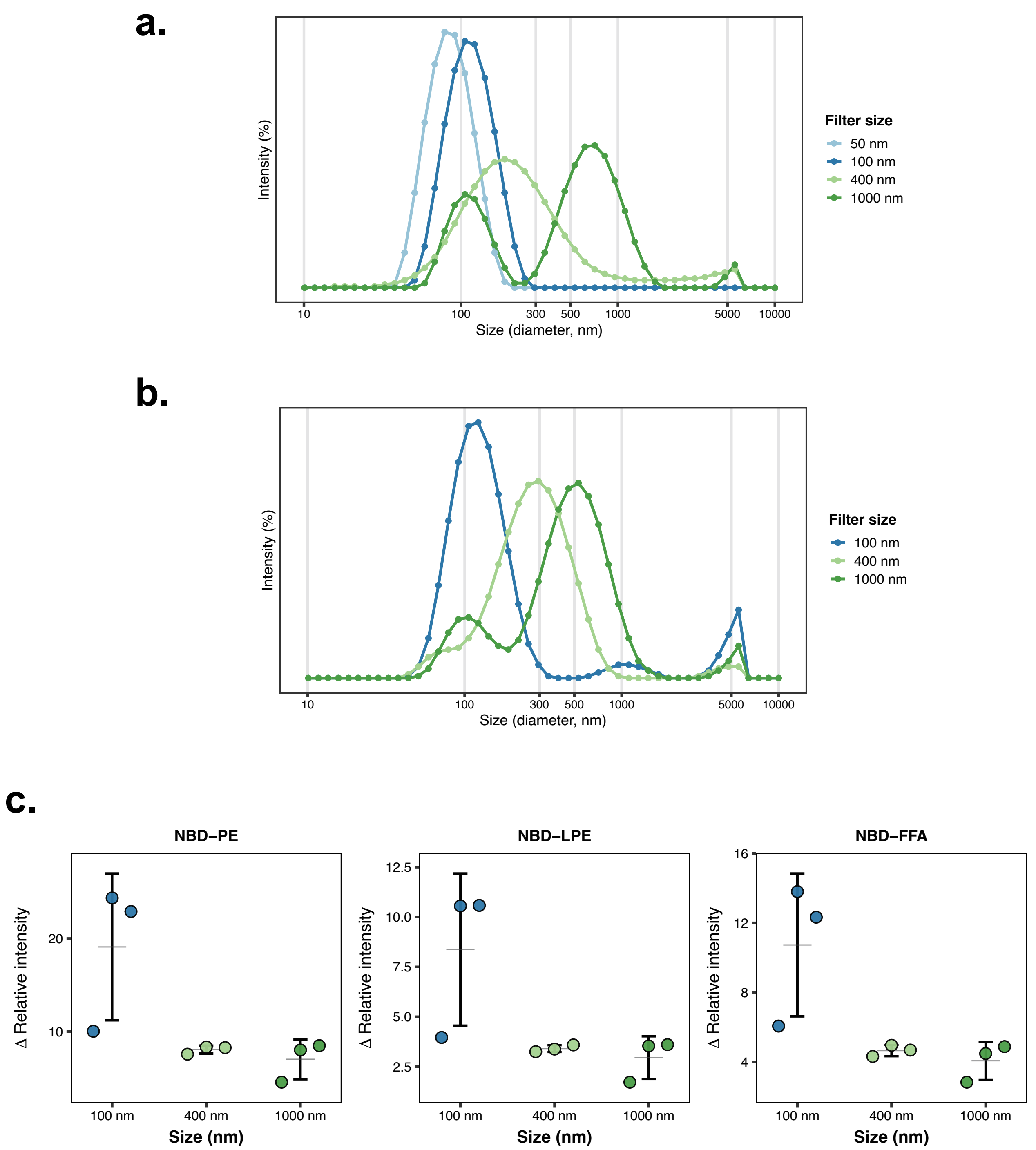
a.
Intensity (%)
Size (diameter, nm)
10
100
300
500
1000
5000
10000
Filter size
50 nm
100 nm
400 nm
1000 nm
b.
Intensity (%)
Size (diameter, nm)
Filter size
100 nm
400 nm
1000 nm
c.
NBD–PE
NBD–LPE
NBD–FFA
Δ Relative intensity
Size (nm)
100 nm
400 nm
1000 nm

Figure. S13

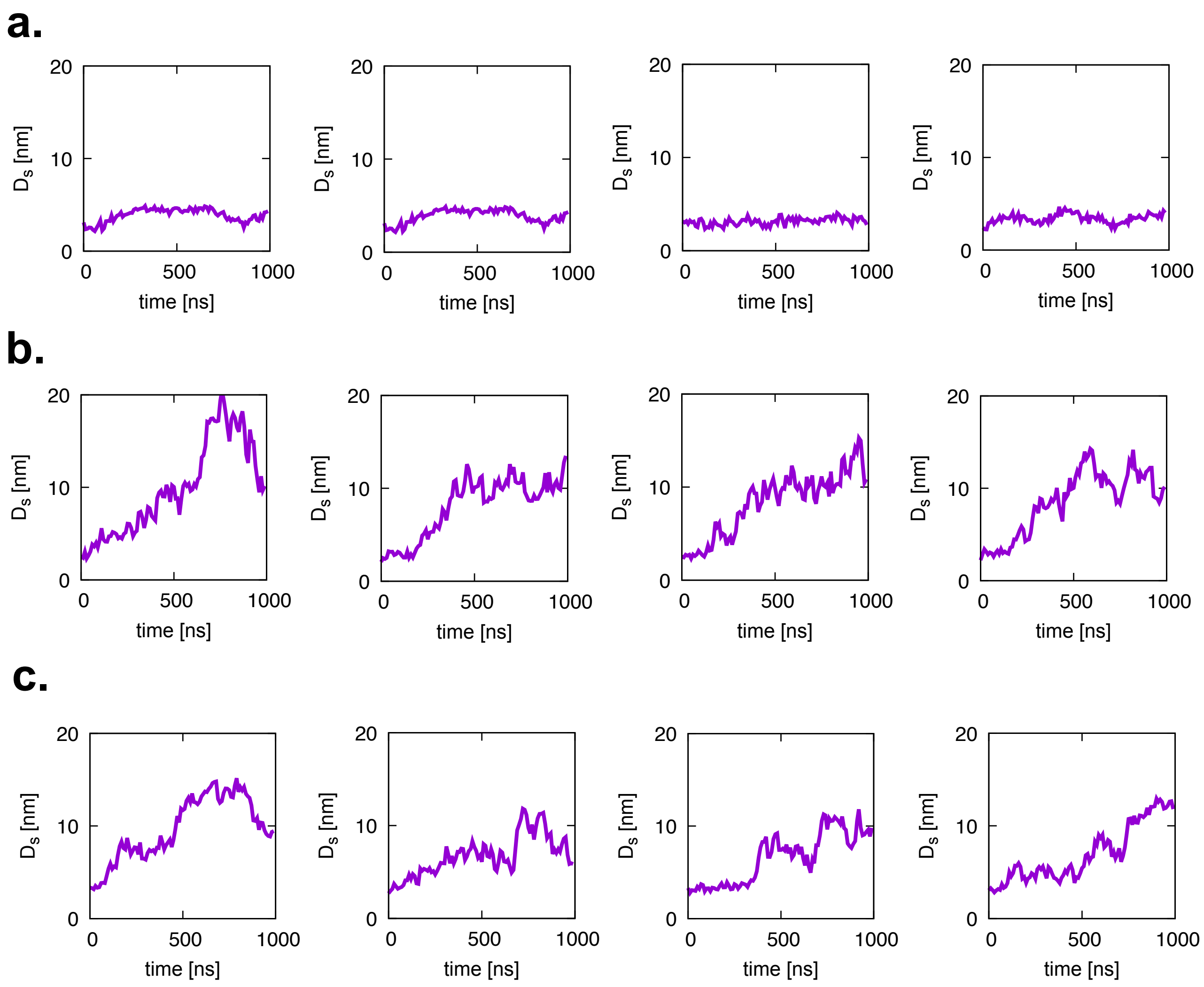


Figure. S14
